# Exploring intra-specific variation in photosynthesis of maize and sorghum

**DOI:** 10.1101/2025.02.21.639328

**Authors:** Rafael L. Almeida, Neidiquele M. Silveira, Heitor L. Sartori, Camila C. Carvalho, Maximiliano S. Scarpari, Aildson P. Duarte, Eduardo Sawazaki, Eduardo C. Machado, Rafael V. Ribeiro

## Abstract

Enhancing crop yield through improved photosynthesis is a key for feeding the global population and providing feedstock for a sustainable green economy. However, effectively linking photosynthetic performance and biomass production in C_4_ species requires an integrative approach at plant canopy. This study aimed to characterize photosynthesis along the canopy of five maize (BM3069-PRO2, AG8701-PRO4, K7500-VIP3, DKB355-PRO3 and B2401-PWU) and sorghum (DKB560, Enforcer, IAC 7021, Brandelisa and Santa Elisa) cultivars, focusing on leaf gas exchange and chlorophyll fluorescence evaluations in three canopy strata: top; middle and bottom. Photosynthetic responses to increasing intercellular CO_2_ concentration and light (*A*–*C*_i_ and *A*–PAR curves, respectively) were performed and key photosynthetic traits estimated. We found a significant variability in the maximum photosynthetic rates across the canopies. Modern maize cultivars exhibited high CO_2_ assimilation in the top and middle canopy leaves, demonstrating physiological adjustments for increasing canopy homogeneity in terms of photosynthesis. Such adjustments included high maximum quantum efficiency of CO_2_ assimilation (ϕ), stomatal conductance, carboxylation rates of PEPC and Rubisco, and leaf nitrogen content (LNC) along the canopy. In contrast, sorghum cultivars showed significant interactions between canopy strata, with DKB560 and Brandelisa standing out for their enhanced CO_2_ uptake and ϕ throughout the plant canopy. Our findings highlight maize as an efficient C_4_ crop, characterized by a high photosynthetic capacity and relative uniform photosynthesis across the canopy. This is attributed to reduced stomatal limitation and higher stomatal conductance, carboxylation of Rubisco (*V*_cmax_) and LNC in the top and middle canopy, along with higher ϕ in middle and bottom layers. Overall, physiological adjustments such as nitrogen redistribution to upper canopy leaves and the optimization of light-use efficiency in lower layers are key for enhancing canopy-level photosynthesis in maize and sorghum cultivars.

## Introduction

Higher crop yield is key for feeding the global population (Ray et al., 2013; Salter et al., 2019) and providing feedstock for a green economy. As we need highly efficient plants in converting sunlight energy into biomass, C_4_ plants play an important role in food and bioenergy supply due to their high photosynthetic efficiency based on the CO_2_ concentration mechanism (CCM). In the mesophyll cells, the phosphoenolpyruvate (PEP) carboxylase (PEPC) fixes the CO_2_ into four-carbon molecule, which is transported to the bundle sheath cells and decarboxylated by a malic enzyme dependent on NADP or NAD (NADP-ME or NAD-ME) and by phosphoenolpyruvate carboxykinase (PEPCK). Such decarboxylation increases the [CO_2_] in bundle sheath cells, where the ribulose-1,5-bisphosphate (RuBP) carboxylase/oxygenase (Rubisco) refixes the CO_2_ into carbohydrates through the C_3_ pathway. Overall, the C_4_ photosynthesis is regulated by the CO_2_ diffusion from the atmosphere to the carboxylation sites, by the carboxylation reactions – PEPC and Rubisco activities – and by the regeneration of RuBP and PEP driven by ATP produced through photochemical reactions (von Caemmerer and Furbank, 2003; Yin et al., 2011; Sales et al., 2018).

Light distribution within plant canopy is an important factor limiting the photosynthetic performance in C_4_ crops (Marchiori et al., 2010, 2014; Jaikumar et al., 2021; Cruz et al., 2022; Almeida et al., 2022). Architecture, size, composition and longevity of the plant canopy affect both quality and quantity of light reaching leaves, with aging and light acclimation along the canopy also changing CO_2_ assimilation (Davey et al., 2017; Almeida et al., 2022). At the canopy scale, plant biomass can be enhanced by changes in plant architecture and higher efficiency of photosynthesis (Marchiori et al., 2010, 2014; Zhu et al. 2010; Slattery et al., 2018). According to Almeida et al. (2022), plant canopy should have three important features for high photosynthesis: (1) top leaves with high photosynthetic rates (high photosynthetic capacity); (2) similar photosynthetic activity in top and bottom leaves in a given light intensity (homogeneous photosynthesis); and (3) bottom leaves photosynthesizing close to the maximum even under low light intensity (light acclimation). These features might boost photosynthesis and then increase yield through breeding, with the selection of top genotypes. However, our knowledge integrating canopy and photosynthesis is limited, mainly in C_4_ species.

C_4_ species present a high range of maximum photosynthetic rates, from 14 to 61 μmol m^−2^ s^−1^, being a possible key trait in breeding programs due its high heritability (Jackson et al., 2016; Li et al., 2017; Almeida et al., 2021). For instance, most of the literature characterizes the C_4_ photosynthetic responses in few genotypes or only in sun-exposed leaves, as done in energy cane (Cruz et al., 2021, 2022), maize and giant *Miscanthus* (Lee et al., 2021), sorghum (Jaikumar et al., 2021) and sugarcane (Marchiori et al., 2010, 2014; Almeida et al., 2021). In addition, no detailed information about how photosynthesis varies along the vertical canopy profile has been provided for modern cultivars of maize and sorghum, two important crops. An integrative perspective considering the canopy profile is needed for linking photosynthetic activity and biomass production in C_4_ species. Here, we aim to characterize photosynthesis along the canopy profile of two ‘NADP-ME type’ C_4_ crop species: maize (*Zea mays* L.) and sorghum (*Sorghum bicolor* L.), revealing the variability within canopy (three leaf ages) and among genotypes (five cultivars) of each species.

## Material and Methods

### Plant growth conditions

Five cultivars of maize (*Zea mays* L.) and sorghum (*Sorghum bicolor* L.) were studied (Table 1). Seeds were sown in plastic pots (12 L) filled with commercial substrate composed of *Sphagnum* spp., rice straw and perlite in 7:2:1 ratio (Carolina Soil of Brazil, Vera Cruz RS, Brazil). The plants were fertilized daily with nutrient solution modified from Sarruge (1975): 15 mmol N L^−1^ (7 % as NH_4_^+^); 4.8 mmol K L^−1^; 5 mmol Ca L^−1^; 2 mmol Mg L^−1^; 1 mmol P L^−1^; 1.2 mmol S L^−1^; 28 μmol B L^−1^; 54 μmol Fe L^−1^; 5.5 μmol Mn L^−1^; 2.1 μmol Zn L^−1^; 1.1 μmol Cu L^−1^; and 0.01 μmol Mo L^−1^. Four plants per genotype of each species were maintained under greenhouse conditions throughout the experimental period, where air temperature averaged 25.5±5.4°C and air relative humidity 79.6±13.1%. To mitigate the potential impact of environmental variations within the greenhouse, the position of the plants was randomized weekly.

**Table 1.**
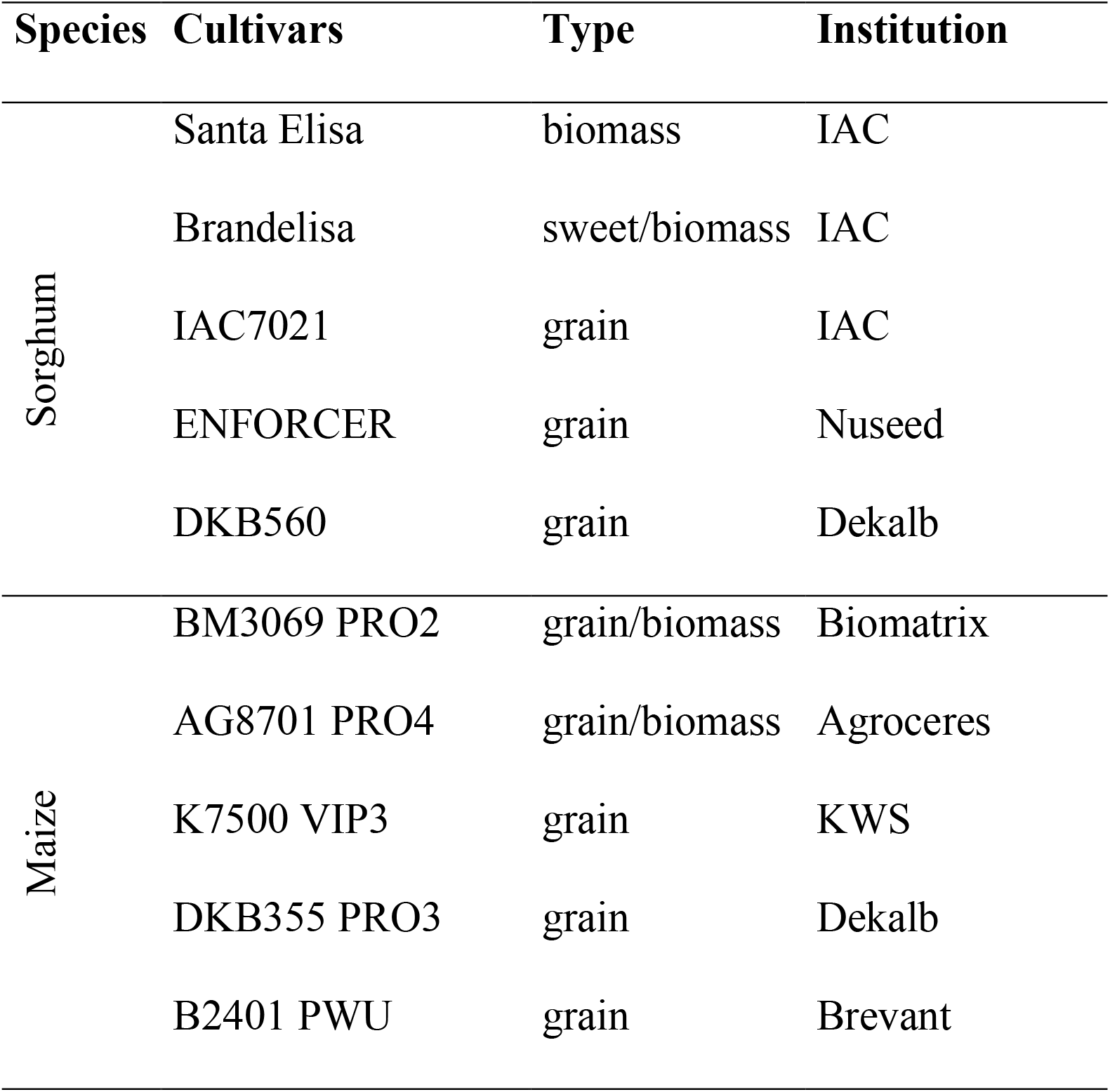
List of maize and sorghum cultivars, type and the institution responsible for breeding.

### Photosynthesis

Leaf gas exchange and chlorophyll fluorescence were evaluated when plants were approximately two months old, corresponding to the R1 stage (Silking) in maize and the stage 6 (Blooming/Flowering) in sorghum. The measurements were taken in leaves from three canopy strata: top; middle; and bottom. For the top canopy, we focused on the flag leaf in maize plants. However, we observed some variations in growth habit among sorghum cultivars. The top canopy was defined by the flag leaf for four sorghum cultivars, except for Santa Elisa. For this cultivar, the top canopy was represented by leaf +1, which is defined as the first fully expanded leaf with visible ligule. Moving to the middle canopy, our attention was on the ear leaf in maize and the third leaf below the flag leaf (or leaf +1) in sorghum. For the bottom canopy, we evaluated the fourth leaf below the ear leaf in maize and the sixth leaf below the flag leaf (or leaf +1) in sorghum, as shown in Fig. S1.

Gas exchange measurements were taken using an infrared gas analyzer IRGA (Li-6400XT, LICOR Inc., Lincoln NE, USA) and a modulated fluorometer (6400-40LCF, LICOR Inc., Lincoln NE, USA) and recorded when the coefficient of variation was less than 2% and there was temporal stability. Measurements of leaf CO_2_ assimilation (*A*), stomatal conductance (g_s_) and the intercellular CO_2_ concentration (*C*_i_) were taken under air temperature of 28°C, with a water vapor pressure deficit lower than 1.5 kPa and varying the photosynthetically active radiation (PAR) and air CO_2_ concentration.

Chlorophyll fluorescence was evaluated using signals emitted before and after a saturation pulse (λ<710 nm; PAR ∼ 8,000 μmol m^−2^ s^−1^; 0.8 s) and after the excitation of the photosystem I (PSI) by far-red light (λ=735 nm; PAR ∼ 5 μmol m^−2^ s^−1^; 3.0 s). Firstly, the leaves were acclimated in the dark for 30 minutes. Mitochondrial respiration in dark (R_d_), the maximum (*F*_m_) and minimum (*F*_o_) fluorescence were measured and the variable fluorescence in dark (*F*_v_*=F*_m_*–F*_o_) and the maximum quantum efficiency of PSII (*F*_v_/*F*_m_) estimated. During leaf gas exchange, we monitored the steady-state (*F*’), maximum (*F*_m_’) and minimum (*F*_o_’, after excitation of the PSI) fluorescence (Baker, 2008). From these data, the effective quantum efficiency of PSII [Φ_PSII_*=*(*F*_m_’-*F*’)/*F*_m_’] and the non-photochemical quenching [NPQ=(*F*_m_-*F*_m_’)/*F*_m_’] were calculated. The relative excess of energy (EXC) was estimated as: EXC=[(*F*_v_*/F*_m_– Φ_PSII_)/(*F*_v_*/F*_m_)], according to Bilger et al. (1995).

The response of photosynthesis to increasing intercellular CO_2_ concentration (*A*–*C*_i_ curve) was evaluated after leaf acclimation (15 minutes) to air CO_2_ concentration (*C*_a_) of 400 μmol mol^−1^ and PAR of 2000 μmol m^−2^ s^−1^. After this time, *C*_a_ inside the cuvette was changed as follows: 400, 300, 200, 120, 85, 70, 55, 400, 400, 500, 600, 800, 1200, and 1500 μmol mol^−1^, with each step taking 3 minutes or coefficient of variation below 2%. The *A*–*C*_i_ curves were fitted (r^2^>0.95) using the equation 1, according to Almeida et al. (2021):

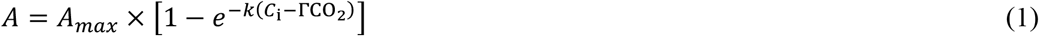

where *A*_*max*_ is the maximum leaf CO_2_ assimilation under high light and CO_2_ saturation, ΓCO_2_ is the CO_2_ compensation point, and *k* is the fitting coefficient.

The apparent maximum carboxylation rates of phosphoenolpyruvate carboxylase (PEPC, *V*_pmax_) and ribulose-1,5-bisphosphate carboxylase/oxygenase (Rubisco, *V*_cmax_) were estimated from the fitted *A*–*C*_i_ curves (equation 1), following Yin et al. (2011):

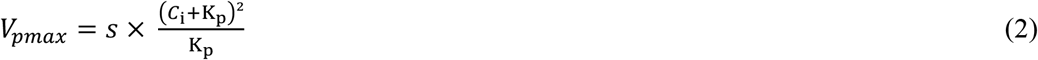

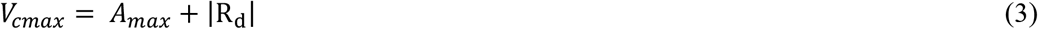

where *s* is the angular coefficient obtained by the initial linear part of *A*–*C*_i_ curves (*C*_i_ < 100 μmol mol^−1^) and K_p_ is the Michaelis-Menten constant of PEPC for CO_2_ (Yin et al., 2016; von Caemmerer, 2021):

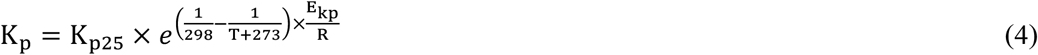

where K_p25_ is Michaelis–Menten constant of PEPC for CO_2_ at 25°C (82 μbar), T is the leaf temperature (°C) during measurements, E_kp_ is the K_p_ activation energy (38.3 kJ mol^−1^), and R is the universal gas constant (0.008314 kJ K^−1^ mol^−1^).

The stomatal limitation of photosynthesis (*L*_S_) was estimated from *A*–*C*_i_ curve, following Long and Bernacchi (2003):

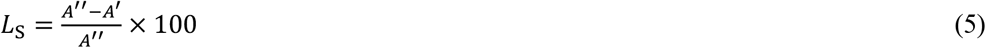

where *A*’’ and *A*’ are leaf CO_2_ assimilation at *C*_i_ and *C*_a_ of 400 μmol mol^−1^, respectively.

The metabolic limitation (*L*_M_) of photosynthesis in plant canopy was also calculated, adapted from Lawlor (2002):

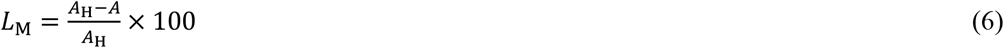

where *A*_H_ is the maximum and *A* is the minimum leaf CO_2_ assimilation at *C*_a_ of 400 μmol mol^−1^ along the canopy profile, considering only rates measured under high light and CO_2_ saturation (at the curve plateau).

The response of photosynthesis to light (*A*–PAR curve) was evaluated after leaf acclimation to light and CO_2_, as done for the *A*–*C*_i_ curves. After leaf gas exchange reached the steady state, PAR inside the Li-6400XT cuvette was changed in the following sequence: 2000, 1500, 1000, 500, 300, 200, 100, 50 and 0 μmol m^−2^ s^−1^, with each step taking 3 minutes or coefficient of variation below 2%. The *A*–PAR curves were fitted using the equation 7, adapted from Marshall and Biscoe (1980):

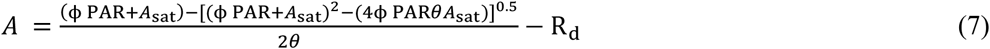

where *A*_sat_ is the maximum CO_2_ assimilation on leaf area basis; ϕ is the maximum quantum efficiency of CO_2_ assimilation; R_d_ is the dark respiration (when PAR = 0 μmol m^−2^ s^−1^); and θ is the curvature factor. From *A*–PAR curves, we also estimated the light compensation point (LCPT, PAR when *A* = 0 μmol m^−2^ s^−1^, from the *x*-axis intercept), according to Heberling and Fridley (2013).

### Leaf area, chlorophyll index and leaf nitrogen content

After leaf gas exchange evaluations, leaf area (LA) was measured using a planimeter model Li-3000C (LICOR Inc., Lincoln NE, USA). Leaf dry matter (DML) was determined after drying all leaves in an oven at 60°C with forced-air circulation until a constant weight was achieved. From these data, the specific leaf mass (SLM = DML/LA, in kg m^−2^) was estimated. The chlorophyll meter model CFL1030 (Falker, Porto Alegre RS, Brazil) was used to evaluate the chlorophyll *a, b, a*/*b* and *a* + *b* (total) indexes. Leaves used for photosynthetic evaluations were collected and their nitrogen concentration (LNC) was determined by the Kjedahl method (Bataglia et al., 1983).

### Experimental design and data analysis

The experimental design was a randomized 3×5 factorial arrangement, with four biological replicates: three canopy levels (top, middle and bottom); and five genotypes for each of the two species studied (maize and sorghum). Normal distribution and homogeneity of variances were tested using the Shapiro-Wilk and Hartley tests, respectively. Following these preliminary tests, all variables were subjected to the analysis of variance (ANOVA). Mean values were compared by the Tukey test (*p*<0.05) when statistical significance was detected. Additionally, the correlation between the photosynthetic traits was analyzed through Pearson’s coefficient. All analyses were conducted using the R software (R Core Team 2024; version 4.4.0, R-project, packages ‘corrplot’, ‘ExpDes’, ‘Hmisc’ and ‘Readxl’).

## Results

### Leaf gas exchange

*A*–*C*_i_ and *A*–PAR curves revealed significant variation in the maximum photosynthetic rates across the canopies of maize (from 22.5 to 50.4 μmol m^−2^ s^−1^) and sorghum (from 11.2 to 46.0 μmol m^−2^ s^−1^, Figs. 1 and S2). To further analyze these findings, we estimated the photosynthetic canopy homogeneity index (CHI) – the ratio of photosynthesis between the top and bottom leaves of each species under high light and high air CO_2_ concentration. For maize, the CHI varied between 1.04 (BM2401) and 1.44 (AG8701). In sorghum, the CHI ranged from 1.19 (DKB560) and 2.16 (Santa Elisa). In terms of metabolic limitation (*L*_M_) of photosynthesis along the canopy profile, we observed differences among the genotypes of both species. *L*_M_ varied from 12.9% (B2401) to 30.9% (AG8701) in maize, whereas it ranged from 16.1% (DKB560) to 53.8% (Santa Elisa) in sorghum, as shown in Fig. 1.

**Fig. 1.**
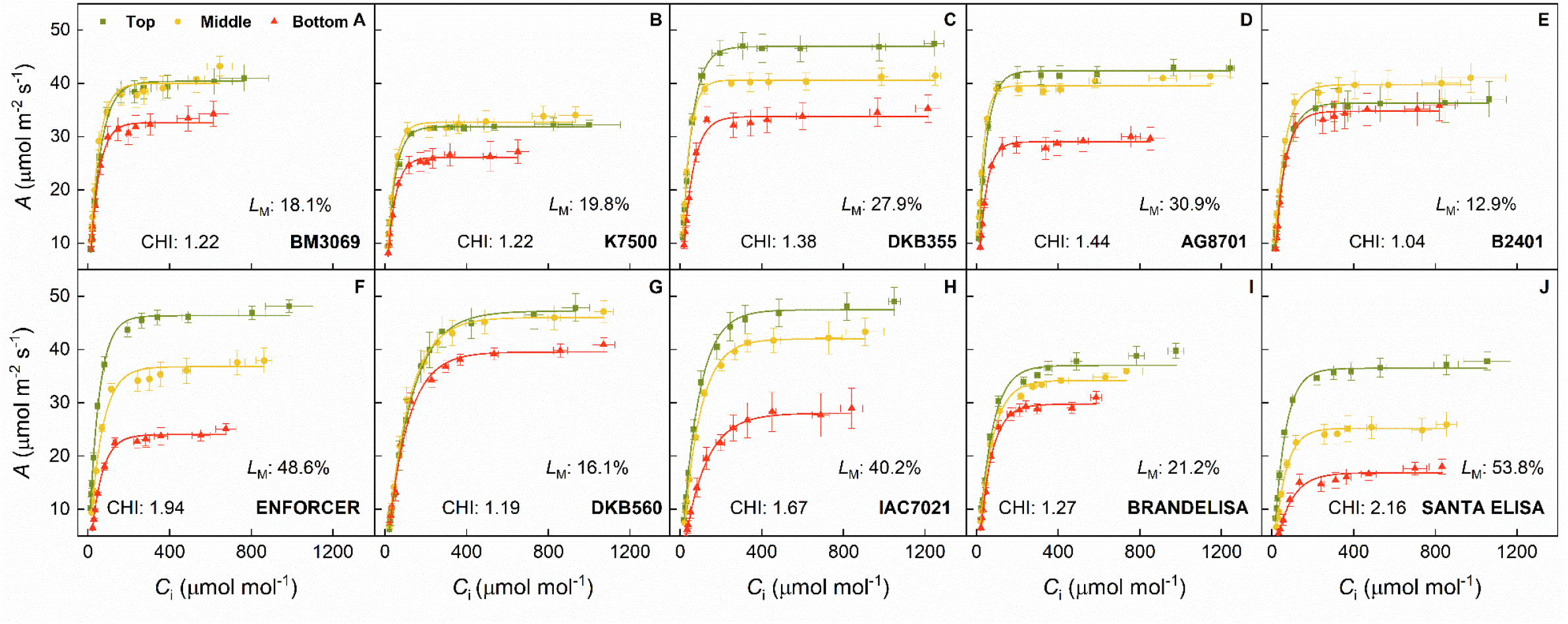
Response curves of leaf CO_2_ assimilation rate (*A*) to intercellular CO_2_ concentration (*C*_i_) in top (green), middle (yellow) and bottom (red) leaves of maize (A-E) and sorghum (F-J) cultivars, with their respective canopy homogeneity index (CHI) and metabolic limitation (*L*_M_). Each symbol represents the mean±standard error (*n*=4).

Our study revealed no significant interaction (*p*>0.05) between the canopy position and the maize cultivars for photosynthesis (*A*_400_) and stomatal conductance (*g*_s400_) measured at *C*_a_ = 400 μmol mol^−1^ (Fig. 2), saturated CO_2_ assimilation (*A*_sat_), the maximum quantum efficiency of CO_2_ assimilation (ϕ) and light compensation point (LCPT; Fig S3). When comparing maize cultivars, K7500 exhibited the lowest *A*_400_, *A*_sat_ and ϕ (Figs. 2A and S3A,C). Generally, the bottom leaves displayed lower *A*_400_ and *A*_sat_ (30.1±1.1 and 39.9±1.8 μmol m^−2^ s^−1^) compared to the top (38.5±1.4 and 56.1±1.9 μmol m^−2^ s^−1^) and middle leaves (37.3±1.0 and 53.7±1.6 μmol m^−2^ s^−1^; Figs. 2B and S3B). Interestingly, this pattern was reversed for ϕ. The middle and bottom leaves had higher ϕ (0.065±0.003 and 0.067±0.003 μmol μmol^−1^) than the top leaf (0.056±0.003 μmol μmol^−1^; Fig. S3D). For stomatal conductance, DKB355 exhibited the highest *g*_s400_ (0.484±0.028 mol m^−2^ s^−1^) among the maize cultivars (Fig. 2C). In general, bottom leaves presented the lowest *g*_s400_ when compared to the middle and top leaves (Fig. 2D). We found no interactions and differences among the genotypes and canopy layers for *L*_S_, which averaged 3.1±0.3%. Regarding the LCPT, K7500 had the highest value (62±5 μmol m^−2^ s^−1^) among maize cultivars (Fig. S3E) and it was lower in bottom leaves (44±2 μmol m^−2^ s^−1^) in relation to the middle and top leaves (Fig. S3F).

**Fig. 2.**
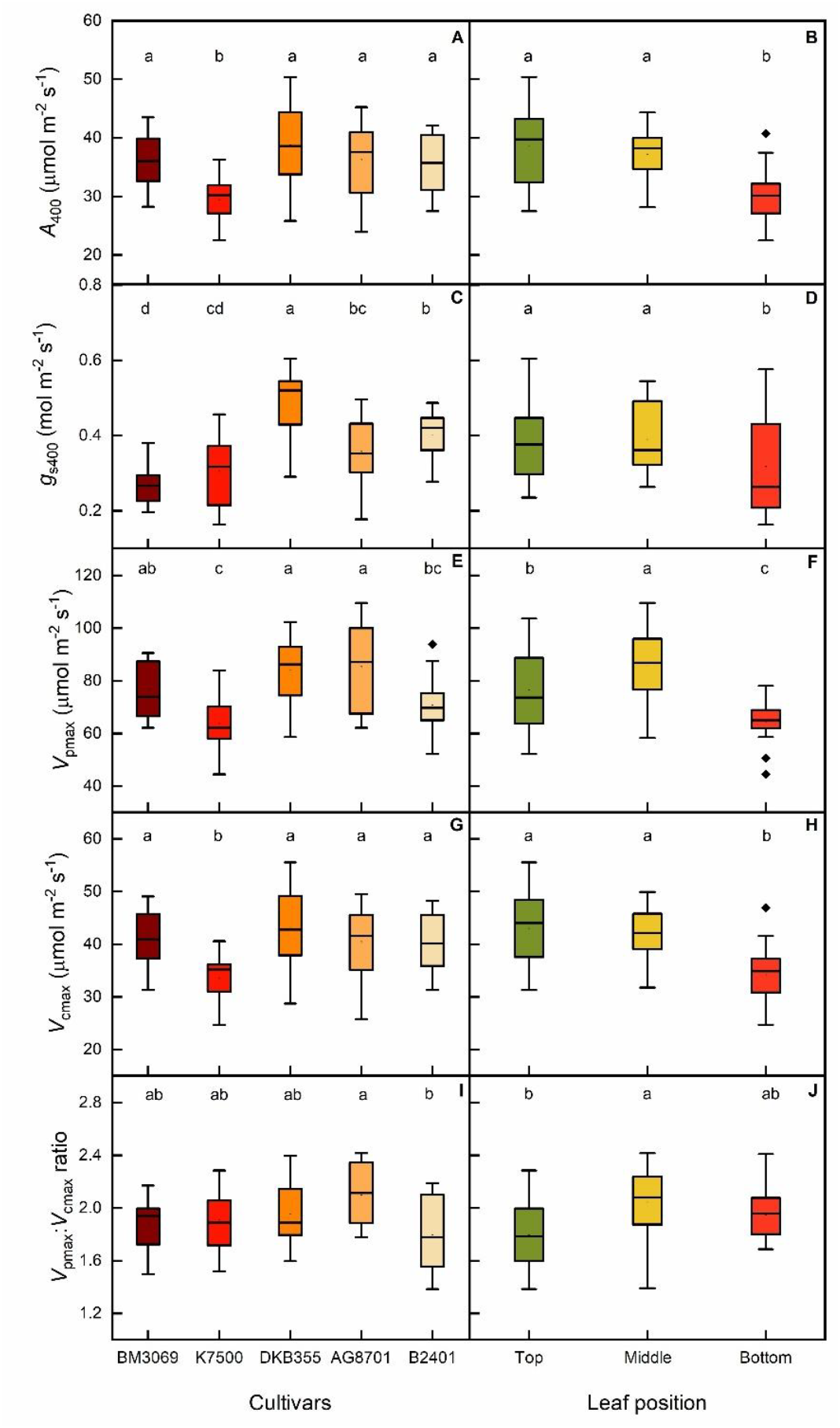
Leaf CO_2_ assimilation (*A*_400_, A and B) and stomatal conductance (*g*_s400_, C and D) at partial air CO_2_ pressure (*C*_a_) of 400 μmol mol^−1^, *in vivo* maximum carboxylation rates of PEPC (*V*_pmax_, E and F) and Rubisco (*V*_cmax_, G and H) and *V*_pmax_:*V*_cmax_ ratio (I and J) in five maize cultivars and three canopy layers. Different letters indicate statistical differences among cultivars (in A, C, E and G, Tukey *p*<0.05, *n*=12), and canopy layers (in B, D, F and H, Tukey *p*<0.05, *n*=20).

Contrary to our findings in maize, there was significant interaction (*p*<0.01) between the canopy layers and the sorghum cultivars for photosynthetic parameters such as *A*_400_, *g*_s400_, *L*_S_, *A*_sat_ and ϕ. Enforcer and Santa Elisa presented the largest variation in *A*_400_ and *A*_sat_ along the canopy. Enforcer, DKB560 and IAC7021 emerged as the superior genotypes for photosynthesis in top leaves, achieving *A*_400_ close to 46 μmol m^−2^ s^−1^ and *A*_sat_ about 55 μmol m^−2^ s^−1^ (Figs. 3A and S4A). Enforcer stood out as the genotype showing the highest ϕ in bottom leaves, with values about 0.077±0.004 μmol μmol^−1^ (Fig. S4B). For stomatal conductance, DKB560 showed no differences along the canopy profile (averaging 0.335±0.014 mol m^−2^s^−1^), while Enforcer, IAC7021, Brandelisa and Santa Elisa exhibited lower *g*_s400_ in the bottom leaves (Fig. 3B). The stomatal limitation (*L*_S_) was higher in the top (19.2±3.2%) and middle (16.0±1.6%) than in the bottom (11.5±1.9%) leaves of DKB560. Conversely, Santa Elisa presented higher *L*_S_ in the bottom (12.4±2.6%) compared to the top and middle leaves (4.8±0.6 and 3.7±1.0%). Enforcer, IAC7021 and Brandelisa showed no differences in *L*_S_ throughout the canopy, averaging 5.3±0.5%, 12.0±1.3% and 8.1±0.8%, respectively (Fig. 4A). No interaction (*p*>0.05) between cultivars and canopy layers was found for light compensation point, with DKB560 and IAC7021 presenting the highest LCPT values (about 54 μmol m^−2^ s^−1^; Fig. S5A). In general, the bottom leaves had LCPT of 39±2 μmol m^−2^ s^−1^, which was about 19.0% and 31.3% lower than those of the middle and top leaves, respectively (Fig. S5B).

**Fig. 3.**
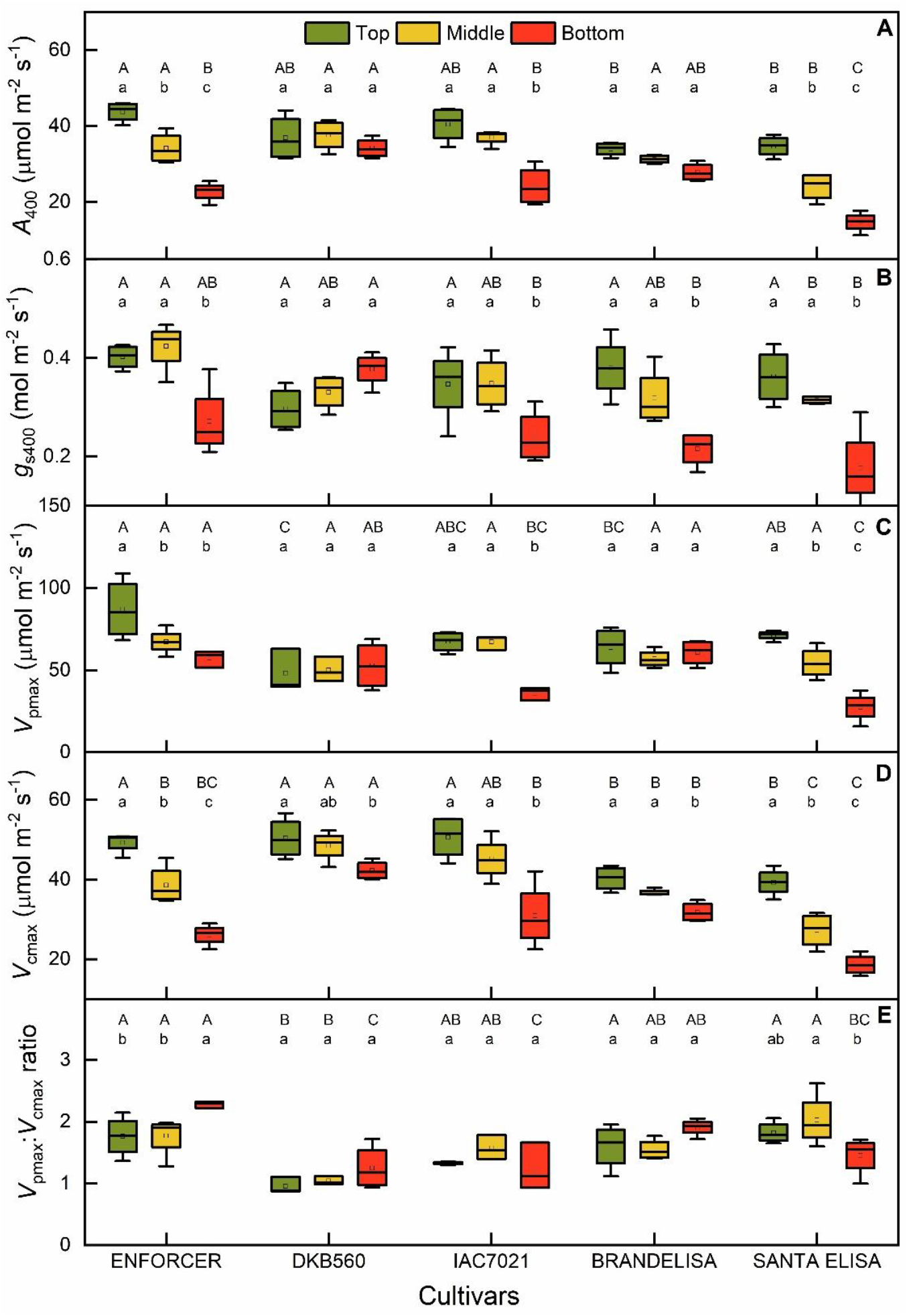
Leaf CO_2_ assimilation (*A*_400_, in A) and stomatal conductance (*g*_s400_, in B) at partial air CO_2_ pressure (*C*_a_) of 400 μmol mol^−1^, *in vivo* maximum carboxylation rates of PEPC (*V*_pmax_, C) and Rubisco (*V*_cmax_, D) and *V*_pmax_:*V*_cmax_ ratio (E) in top (green), middle (yellow) and bottom (red) leaves of five sorghum cultivars. Different capital and lowercase letters indicate statistical differences among cultivars and canopy layers, respectively (Tukey *p*<0.05, *n*=4).

**Fig. 4.**
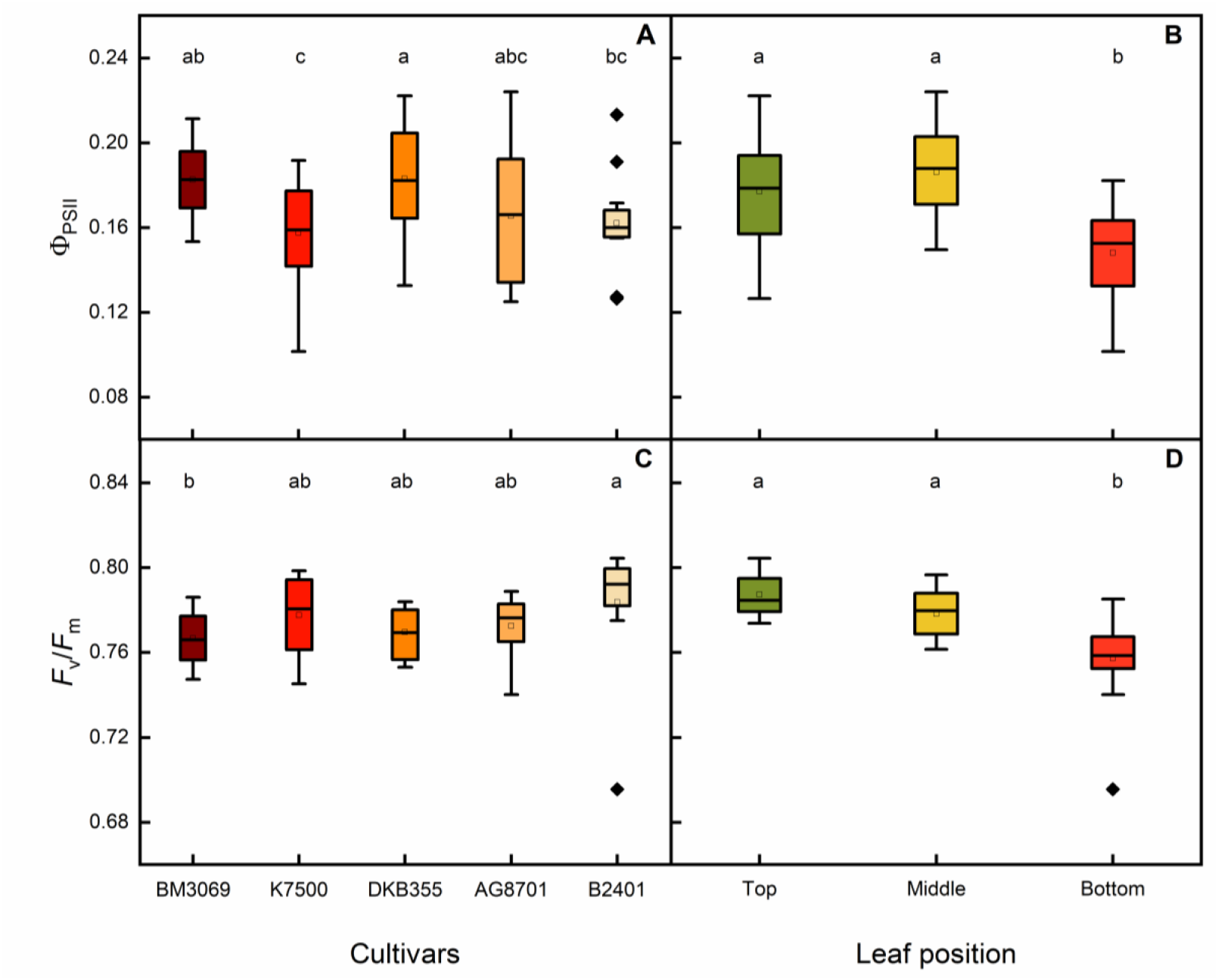
Effective (Φ_PSII_, A and B) and maximum (*F*_v_/*F*_m_, C and D) quantum efficiency of PSII in five maize cultivars and three canopy layers. Different letters indicate statistical differences among cultivars (A and C, Tukey *p*<0.05, *n*=12) and canopy layers (B and D, Tukey *p*<0.05, *n*=20).

Mitochondrial dark respiration (*R*_d_) did not change among the maize cultivars (*p*>0.05), with an average of 3.3±0.1 μmol m^−2^ s^−1^. However, differences were noticed among canopy strata. The top and middle maize leaves (3.6±0.2 and 3.5±0.1 μmol m^−2^ s^−1^, respectively) presented higher *R*_d_ than bottom leaves (2.9±0.1 μmol m^−2^ s^−1^). In sorghum, we found no interaction between genotypes and the canopy strata. Herein, DKB560 and IAC7021 presented the highest *R*_d_ (about 2.7 μmol m^−2^ s^−1^), and top leaves had higher *R*_d_ (2.9±0.1 μmol m^−2^ s^−1^) when compared to the middle and bottom leaves (2.2±0.1 to 2.4±0.1 μmol m^−2^ s^−1^, respectively).

### Biochemistry and photochemistry

No interaction (*p>*0.05) between the canopy strata and maize cultivars was found for *V*_pmax_, *V*_cmax_, *V*_pmax_:*V*_cmax_ ratio, Φ_PSII_ and *F*_v_*/F*_m_. BM3069, AG8701 and DKB355 presented higher *V*_pmax_ than K7500 and B2401 (Fig. 2E), whereas K7500 presented the lowest *V*_cmax_ among all maize cultivars (Fig. 2G). The *V*_pmax_:*V*_cmax_ ratio ranged from 1.8 to 2.1, with AG8701 presented higher values than B2401 (Fig. 2I). When comparing canopy strata, both *V*_pmax_ and *V*_cmax_ were lower in the bottom one (Fig. 2F, H), while the lower *V*_pmax_:*V*_cmax_ ratio was found in the top leaves (Fig. 2J). Φ_PSII_ and *F*_v_*/F*_m_ also varied among the genotypes, with DKB355 and B2401 exhibiting the highest Φ_PSII_ and *F*_v_*/F*_m_ values, respectively (Fig. 5A,C). In general, the bottom leaves showed the lowest Φ_PSII_ (0.148±0.004) and *F*_v_*/F*_m_ (0.757±0.004) when compared to middle (0.186±0.005 and 0.778±0.002) and top (0.177±0.006 and 0.787±0.002) leaves (Fig. 5B,D).

**Fig. 5.**
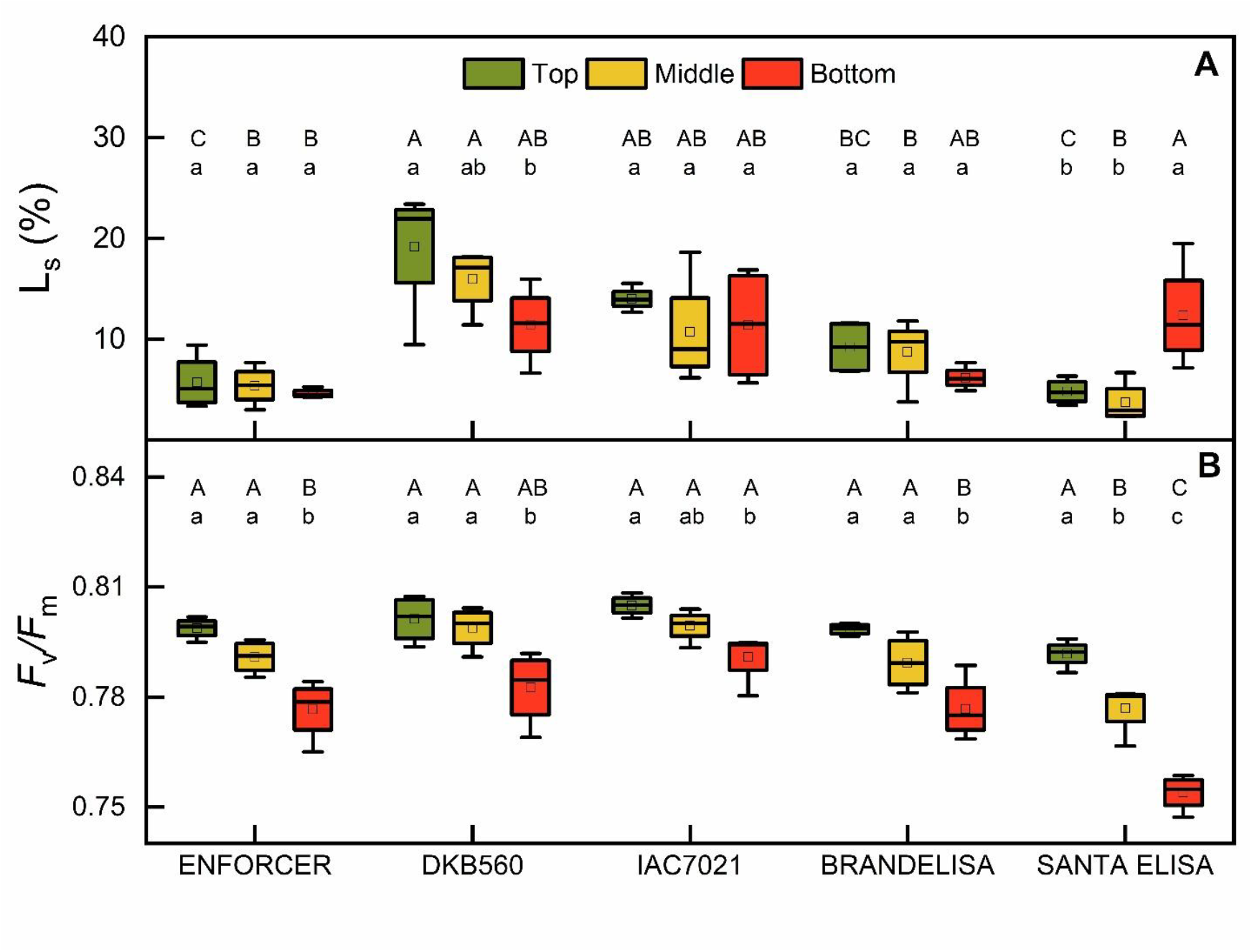
Stomatal limitation of photosynthesis (*L*_s_, in A) and maximum quantum efficiency of PSII (*F*_v_*/F*_m_, in B) in top (green), middle (yellow) and bottom (red) leaves of five sorghum cultivars. Different capital and lowercase letters indicate statistical differences among cultivars and canopy layers, respectively (Tukey *p*<0.05, *n*=4).

In sorghum species, we found significant interaction (*p<*0.05) between the canopy layers and cultivars for *V*_pmax_, *V*_cmax_, *V*_pmax_:*V*_cmax_ ratio and *F*_v_*/F*_m_. Nonetheless, no interaction (*p>*0.05) was observed for Φ_PSII_. Enforcer, IAC7021 and Santa Elisa decreased *V*_pmax_ from top to the bottom canopy, contrasting to DKB560 and Brandelisa. Regarding *V*_cmax_, all genotypes presented higher values in the top when compared to the bottom leaves (Fig. 3C,D), while DKB560, IAC7021 and Brandelisa showed no differences for *V*_pmax_:*V*_cmax_ ratio throughout the canopy (Fig. 3E). Regarding Φ_PSII_, no variation was found along canopy profile, which averaged 0.157±0.005. However, the top (0.180±0.008) and middle (0.165±0.009) leaves were more efficient than the bottom (0.127±0.009) ones. The lowest *F*_v_*/F*_m_ was found in the bottom leaves, with Santa Elisa showing the most significant reduction between the top and bottom (Fig. 4B).

### Nitrogen content, leaf area and chlorophyll index

No interaction (*p>*0.05) was observed between canopy strata and maize or sorghum cultivars for LNC. In maize, DKB355, AG8701 and B2401 exhibited higher LNC than BM3069 (Fig. 6A). Among sorghum cultivars, the grain-types Enforcer, DKB560 and IAC7021 showed greater LNC than Brandelisa and Santa Elisa (Fig. 6C). Across both species, LNC was highest in the top and middle canopy strata (Fig. 6B,D).

**Fig. 6.**
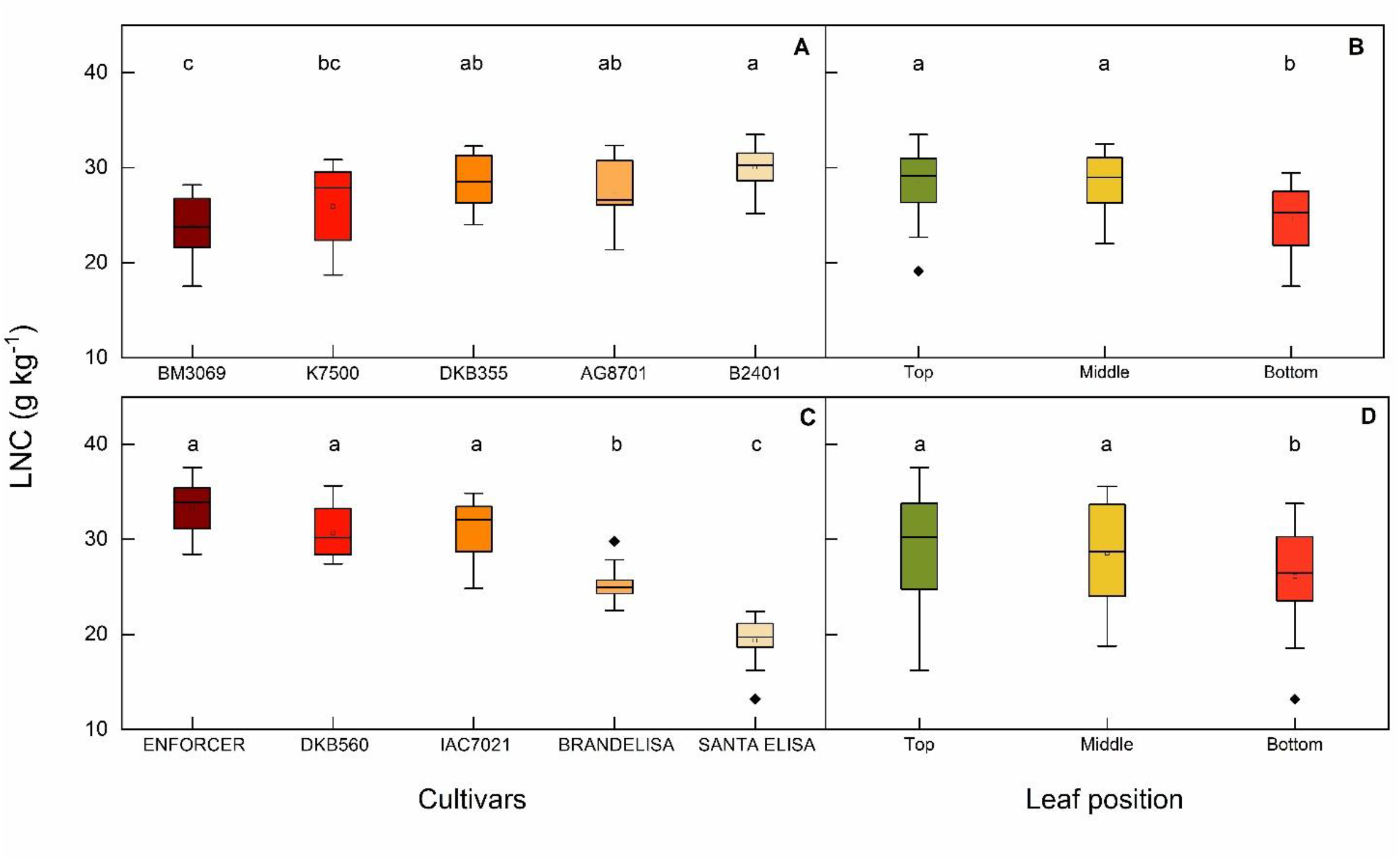
Leaf nitrogen content (LNC) in five maize (A and B) and sorghum (C and D) cultivars and three canopy layers (B and D). Different letters indicate statistical differences among cultivars (A and C, Tukey *p*<0.05, *n*=12) and canopy layers (B and D, Tukey *p*<0.05, *n*=20).

Among maize cultivars, K7500 presented the largest leaf area (1.01±0.03 m^2^) and the lowest specific leaf mass (SLM, ∼44 g m^−2^) when plants were about two months old (Fig. 7A,C). When considering sorghum cultivars, Santa Elisa had the largest leaf area (1.74±0.05 m^2^), while Enforcer had the lowest SLM (∼26 g m^−2^, Fig. 7B,D). No interaction (*p*>0.05) between the canopy strata and maize cultivars was found for chlorophyll *a, b, a*/*b* ratio and total (*a*+*b*). There was no difference in chlorophyll *a* among maize genotypes. DKB355 presented higher chlorophyll *b* (20.5±1.0 *vs*. 15.2±0.7) and total (65.5±1.2 *vs*. 59.0±1.2); and lower *a*/*b* ratio (2.25±0.11 *vs* 2.93±0.12) when contrasted to B2401, as shown in Fig. S6A,C,E,G. Chlorophyll *a* remained uniform across the top, middle and bottom maize leaves. Contrarily, chlorophyll *b* and the total (*a*+*b*) chlorophyll were higher in middle as compared to top leaves, while *a*/*b* ratio was higher in the top ones (Fig. S6B,D,F,H).

**Fig. 7.**
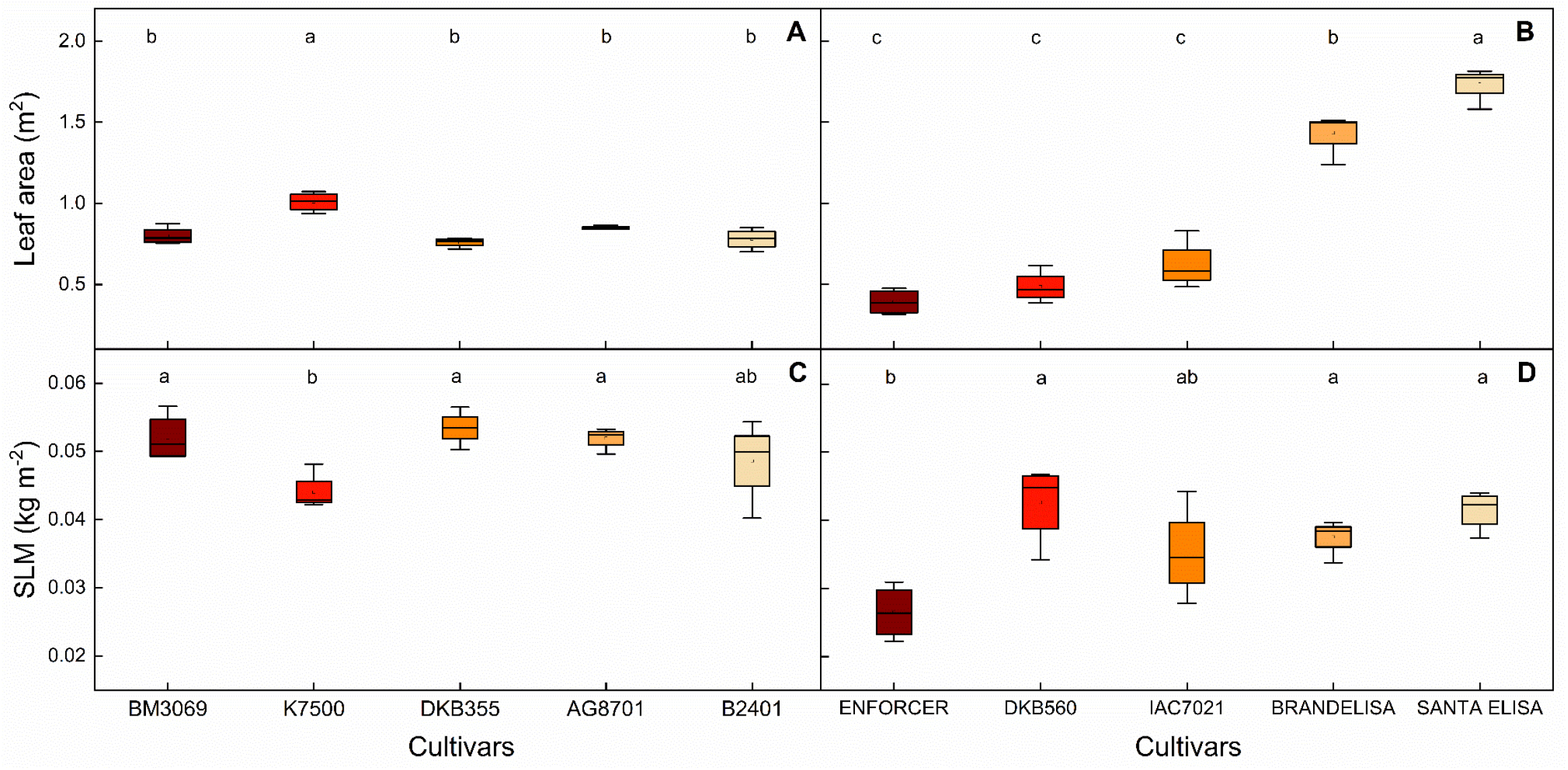
Leaf area (A and B) and the specific leaf mass (SLM, in C and D) in five maize (A and C) and sorghum (B and D) cultivars. Different letters indicate significant differences among cultivars of each species (Tukey, *p*<0.05, *n*=4).

In sorghum, DKB560 showed higher chlorophyll *a* than IAC7021 (Fig. S7A). However, chlorophyll *a* did not show any significant variation (*p*>0.05) across canopy strata, averaging 40.3±0.2 (Fig. S7B). There were significant interactions (*p*<0.01) between the canopy strata and sorghum cultivars when considering chlorophyll *b, a*/*b* ratio and total (*a*+*b*). The top leaves of Enforcer had higher chlorophyll *b* and total than the bottom leaves, while the opposite was true for the *a*/*b* ratio. Furthermore, IAC7021 differed from Enforcer for chlorophyll *b, a*/*b* ratio and total in both top and bottom leaves (Fig. S8). Unfortunately, chlorophyll data of Brandelisa and Santa Elisa cultivars are missing.

### Correlations

*A*_400_ was correlated with *g*_s400_ (r=0.61 and 0.78), *V*_pmax_ (r=0.84 and 0.73), *V*_cmax_ (r=0.998, and 0.97), ϕ_PSII_ (r=0.87 and 0.94), NPQ (r=−0.82 and −0.73), EXC (r=−0.84 and −0.93) and LNC (r=0.66 and 0.67) across maize and sorghum canopies. Notably, both species exhibited a positive correlation between LNC and *V*_cmax_ (r=0.65 and 0.69), as well as between LNC and *F*_v_/*F*_m_ (r= 0.70 and 0.67). Additionally, NPQ and EXC were well correlated in maize (*r*=0.75) and sorghum (*r*=0.81) canopies (Figs. S9 and S10).

## Discussion

### Variation of photosynthesis along the maize canopy profile: physiological adjustments for increasing homogeneity

Our data revealed physiological adjustments and no variation in photosynthetic traits when comparing the top and middle canopy leaves in maize plants. The decline in *A*_400_ and *V*_cmax_ in the bottom leaves are potentially linked to aging and decreases in leaf nitrogen content (LNC; Fig. 6B). In this context, we believe that the primary N source for remobilization in bottom leaves might be the protein Rubisco, as evidenced by the strong relationship of LNC with *A*_400_, *V*_cmax_ and *F*_v_/*F*_m_ (Fig. S9). This process would redistribute nitrogen to upper leaves, sustaining their ability to perform high photosynthetic rates (Yin et al., 2011; Niinemets et al., 2015; Jaikumar et al., 2021). Interestingly, we found an increase in *V*_pmax_:*V*_cmax_ ratio and ϕ in the middle and bottom leaves (Fig. 4J). According to Hikosaka and Terashima, (1995), N remobilization did not decrease ϕ when light is not a limiting factor, as found in our study and in giant *Miscanthus* (Pignon et al., 2017). This observation suggests the photosynthetic acclimation along the maize canopy to optimize the photosynthetic capacity in the top and middle leaves, as found in sugarcane (Almeida et al., 2022). Simultaneously, such acclimation enhances the quantum efficiency of CO_2_ assimilation in the bottom leaves which are several weeks older than top and middle leaves. Decreases in LCPT were also observed in the bottom leaves (Fig. S3), suggesting again a physiological strategy to optimize light-use efficiency.

*V*_pmax_ is a physiological index associated with the capacity of PEPC and subsequently with the ability to increase [CO_2_] in bundle-sheath cells (Bailey et al., 2000). High activities and abundance of PEPC, Rubisco, pyruvate orthophosphate dikinase (PPDK), phosphoenolpyruvate carboxykinase (PEPCK) and NADP-dependent malate dehydrogenase are crucial for maintaining photosynthesis within the deeper canopy, where leaves are shaded (Vu et al., 2006; Sales et al., 2018, Jaikumar et al., 2021). The efficiency of the CO_2_ concentrating mechanism (CCM) plays a significant role in this context. If CCM is effective, photorespiration is inhibited, leakiness is low and the quantum efficiency of CO_2_ assimilation is high (Farquhar, 1983; Kromdijk et al., 2008; Sales et al., 2018). Particularly, high *V*_pmax_:*V*_cmax_ ratio and a small response (3.0±0.3%) of photosynthesis when changing *C*_a_ from 400 to 1500 μmol mol^−1^ (Figs. 1A-E and 2J) provide compelling evidence of high CO_2_:O_2_ ratio around the active sites of Rubisco in the middle and bottom leaves. Overall, maize photosynthesis has a similar pattern among genotypes and canopy strata, which is likely a consequence of rigorous selection pressure in breeding programs. Such programs might have inadvertently selected genotypes with higher and homogeneous canopy photosynthesis while aiming for higher yield (Lee and Tollernaar, 2007; Andorf et al., 2019; Muntean et al., 2022).

### Underlying factors changing the photosynthetic activity along the sorghum canopy

In contrast to maize, sorghum genotypes exhibited significant variation of photosynthetic traits across the canopy. The source of this variability in sorghum can be traced back to breeding purposes, such as grain yield, sugar accumulation in stalks and biomass/forage production (Table 1, Fig. 11B). Under identical light and [CO_2_] conditions, we identified three distinct patterns of photosynthesis across the canopy of sorghum genotypes: #1-uniform photosynthetic activity across top, middle and bottom leaves, as found in DKB560 and Brandelisa; #2-photosynthetic activity in top and middle leaves that differed from the bottom leaves, as in IAC7021; and #3-photosynthetic activity varying among the top, middle and bottom leaves, as observed in Enforcer and Santa Elisa.

Regarding the pattern #1, DKB560 exhibited high stomatal limitation (*L*_S_) in the top (19.2±3.2%), middle (16.0±1.6%) and bottom (11.5±1.9%) leaves as compared to other cultivars, without any difference in *V*_pmax_ across its canopy. Brandelisa also exhibited a uniform CO_2_ assimilation and *V*_pmax_ throughout the canopy, but with *V*_cmax_ (31.9±1.3 μmol m^−2^ s^−1^) and *L*_S_ (8.1±0.8%) lower than those found in DKB560 (Figs. 3C,D and 4A). However, high *A*_400_ and low metabolic limitation (from 16 to 21%; Fig. 1G,I) found here for DKB560 and Brandelisa indicate an enhanced capacity for CO_2_ fixation across the canopy. Both cultivars presented higher ϕ values in the bottom than the top leaves, a response similar to one found in maize.

When considering the patterns #2 and #3, there was a pronounced reduction in photosynthesis along the canopies of Enforcer, IAC7021 and Santa Elisa. This decrease could be primarily attributed to the N remobilization from bottom leaves to upper ones (Figs. 3A and 6D). Given that a substantial portion leaf N is allocated to photosynthetic enzymes (Sage and Pearcy 1987; Sage 2002), we would speculate that such remobilization may lead to a reduction in both Rubisco abundancy and activity. If this is true, this phenomenon seems to occur without any decrease in ϕ, consistent with previous findings in maize (Hikosaka and Terashima, 1995). In this study, *A*_400_, *V*_cmax_, *F*_v_/*F*_m_ and LNC exhibited remarkably similar patterns across the sorghum canopy (Figs. 3A,C,D and 6C,D), suggesting a strong interrelation among these traits, supported by the correlation analyses (Fig. S10).

### Breeding for enhancing photosynthesis: Is there room to boost canopy efficiency?

While maize genotypes had low variability for photosynthesis and leaf area, sorghum genotypes presented high variability for photosynthetic traits along the canopy profile. Such variation in sorghum indicates the existence of a broad genetic diversity that could be explored to enhance crop yield, as high heritability is found for photosynthesis and leaf area in C_4_ plants (Jackson et al., 2016; Li et al., 2017; Almeida et al., 2021). As several studies have shown a strong link between high photosynthetic activity and yield (Yoon et al., 2020; Ainsworth et al., 2021; De Souza et al., 2022; Wei et al., 2022), photosynthetic traits might be a tool for selecting and screening sorghum genotypes with high yield potential. In maize, the low variability found here is in accordance with the low genetic variation within the maize germplasm, which poses difficulties for enhancing the photosynthetic capacity through breeding (Richards, 2000; Ahmadzadeh et al., 2004).

The biomass-producing sorghum genotypes Santa Elisa and Brandelisa exhibited a large leaf area (Fig. 7B). This phenotype not only enhances the visual appeal of the plant but also suggests a prolonged functional longevity of canopy in terms of CO_2_ uptake and sustained chlorophyll content (Thomas and Howarth, 2000). Notably, Brandelisa exhibited a uniform CO_2_ uptake and *V*_pmax_ throughout the canopy, along with a higher ϕ in the bottom canopy (Figs. 3A and S4B). This pattern might be associated with its ability to maintain chlorophyll content, as observed in DKB560 (Fig. S8). This strategy optimizes the photosynthetic performance in the middle and bottom layers, representing an adaptive mechanism to enhance light-use efficiency (Almeida et al., 2022) with potential to enhance crop productivity. These findings open avenues for future research and breeding programs aimed at enhancing crop productivity in an era of climate change and growing food demand.

## Conclusion

Our study has provided a comprehensive analysis of the variation of photosynthetic traits across maize and sorghum canopies. We highlighted maize as an efficient C_4_ crop, showing high photosynthetic capacity and relative uniform photosynthesis throughout the canopy. This is attributed to reduced stomatal limitation and higher stomatal conductance, Rubisco activity (*V*_cmax_) and LNC in the top and middle canopy, along with higher ϕ in middle and bottom layers. In contrast, sorghum cultivars presented significant variability, with DKB560 and Brandelisa standing out as cultivars with enhanced CO_2_ uptake and ϕ through the canopy. The CO_2_ uptake, ϕ and leaf area variability in sorghum are important traits for crop improvement. Overall, physiological adjustments such as nitrogen redistribution to upper canopy leaves and the optimization of light-use efficiency in lower canopy layers are key for improving canopy photosynthesis in maize and sorghum.

## Acknowledgements

RLA acknowledges the scholarship provided by the Coordination for the Improvement of Higher Education Personnel (Capes, Brazil, Grant n° 88887.646333/2021-00). ECM and RVR are fellows of the National Council for Scientific and Technological Development (CNPq, Brazil, Grants #303904/2023-2 and #304295/2022-1).

## Declarations

No conflict of interests

## Authors’ contributions

RLA, ECM and RVR conceptualized and designed the experiment. APD and ES provided maize and sorghum seeds. RLA, NMS, HLS, MSS and CCC conducted the experiment. RLA and RVR wrote the manuscript, with all authors contributing to data interpretation, discussion and refining the final version of the manuscript.

## Notes

### Competing Interest Statement

The authors have declared no competing interest.

## References

Ahmadzadeh A, Lee EA, Tollenaar M (2004) Heterosis for leaf CER during the grain-filling period in maize. Crop Sci. 44:2095–2100. 10.2135/cropsci2004.2095

Ainsworth EA, Long SP (2021) 30 years of free-air carbon dioxide enrichment (FACE): What have we learned about future crop productivity and its potential for adaptation? Global Change Biology 27:27–49. 10.1111/gcb.15375

Almeida RL, Silveira NM, Miranda MT, Pacheco VS, Cruz LP, Xavier MA, Machado EC, Ribeiro RV (2022) Evidence of photosynthetic acclimation to self-shading in sugarcane canopies. Photosynthetica 60:521–528. 10.32615/ps.2022.045

Almeida RL, Silveira NM, Pacheco VS, Xavier MA, Machado EC, Ribeiro RV (2021) Variability and heritability of photosynthetic traits in Saccharum complex. Theoretical and Experimental Plant Physiology 33:343–355. 10.1007/s40626-021-00217-x

Andorf C, Beavis WD, Hufford M, Smith S, Suza WP, Wang K, Woodhouse M, Yu J, Lübberstedt T (2019) Technological advances in maize breeding: past, present and future. Theor Appl Genet. 132:817–849. 10.1007/s00122-019-03306-3.

Bailey KJ, Battistelli A, Dever LV, Lea PJ, Leegood RC (2000) Control of C4 photosynthesis: effects of reduced activities of phosphoenolpyruvate carboxylase on CO2 assimilation in Amaranthus edulis L. J Exp Bot 51:339–346. 10.1093/jexbot/51.suppl_1.339

Baker NR (2008) Chlorophyll fluorescence: a probe of photosynthesis in vivo. Annual Review in Plant Biology 59:89–113. 10.1146/annurev.arplant.59.032607.092759

Bataglia OC, Furlani AMC, Teixeira JPF, Furlani PR, Gallo JR (1983) Métodos de análise química de plantas. Boletim técnico IAC 78. Instituto Agronômico, Campinas.

Bilger W, Schreiber U, Bock M (1995) Determination of the quantum efficiency of photosystem II and non-photochemical quenching of chlorophyll fluorescence in the field. Oecologia, 102:425–432. 10.1007/BF00341354

Cruz LP, Machado EC, Ribeiro R.V (2022) Estimating the light conversion efficiency by sugarcane: the segmented approach. ? An. Acad. Bras. Cienc. 94:e20211317. 10.1590/0001-3765202220211317

Cruz LP, Pacheco VS, Silva LM, Almeida RL, Miranda MT, Pissolato MD, Machado EC, Ribeiro RV (2021) Morpho-physiological bases of biomass production by energy cane and sugarcane: A comparative study. Ind. Crops Prod. 171:113884. 10.1016/j.indcrop.2021.113884

Davey CL, Jones LE, Squance M, Purdy SJ, Maddison AL, Cunniff J, Donnison I, Clifton-Brown J (2017) Radiation capture and conversion efficiencies of Miscanthus sacchariflorus, M. sinensis and their naturally occurring hybrid M. x giganteus. ? Glob. Change Biol. Bioenergy 9:385–399. 10.1111/gcbb.12331

De Souza AP, Burgess SJ, Doran L, Hansen J, Manukyan L, Maryn N, Gotarkar D, Leonelli L, Niyogi KK, Long SP (2022) Soybean photosynthesis and crop yield are improved by accelerating recovery from photoprotection. Science. 377:851–854. 10.1126/science.adc98

Farquhar GD (1983) On the nature of carbon isotope discrimination in C4 species. Funct Plant Biol 10:205–226. 10.1071/PP9830205

Heberling JM, Fridley JD (2013) Resource-use strategies of native and invasive plants in Eastern North American forests. New Phytologist 200:523–533. 10.1111/nph.12388

Hikosaka K, Terashima I (1995) A model of the acclimation of photosynthesis in the leaves of C3 plants to sun and shade with respect to nitrogen use. Plant Cell & Environment 18:605– 618. 10.1111/j.1365-3040.1995.tb00562.x

Jackson P, Basnayake J, Inman-Bamber G, Lakshmanan P, Natarajan S, Stokes C (2016). Genetic variation in transpiration efficiency and relationships between whole plant and leaf gas exchange measurements in Saccharum spp. and related germplasm. Journal of Experimental Botany 67:861–871. 10.1093%2Fjxb%2Ferv505

Jaikumar NS, Stutz SS, Fernandes SB, Leakey ADB, Bernacchi CJ, Brown PJ, Long SP (2021) Can improved canopy light transmission ameliorate loss of photosynthetic efficiency in the shade? An investigation of natural variation in Sorghum bicolor. Journal of Experimental Botany 72:4965–4980. 10.1093/jxb/erab176

Kromdijk J, Schepers HE, Albanito F, Fitton N, Carrol F, Jones MB, Finnan J, Lanigan GJ, Griffths H (2008) Bundle sheath leakiness and light limitation during C4 leaf and canopy CO2 uptake. Plant Physiol 148:2144–2155. 10.1104/pp.108.129890

Lawlor DW (2002) Limitation to photosynthesis in water-stressed leaves: stomata vs. metabolism and the role of ATP. Ann. Bot. 62:3235–3246. 10.1093/aob/mcf110

Lee EA, Tollenaar M (2007) Physiological basis of successful breeding strategies for maize grain yield. Crop Sci. 47:1–14. 10.2135/cropsci2007.04.0010IPBS

Lee M-S, Boyd RA, Ort DR (2022) The photosynthetic response of C3 and C4 bioenergy grass species to fluctuating light. GCB Bioenergy 14:37–53. 10.2135/cropsci2007.04.0010IPBS

Li C, Jackson P, Lu X, Xu C, Cai Q, Basnayake J, Lakshmanan P, Ghannoum O, Fan Y (2017) Genotypic variation in transpiration efficiency due to differences in photosynthetic capacity among sugarcane related clones. Journal of Experimental Botany 68:2377–2385. 10.1093/jxb/erx107

Long SP, Bernacchi CJ (2003) Gas Exchange measurements, what can they tell us about the underlying limitations of photosynthesis? Procedures and sources of error. Journal of Experimental Botany 54:2393–2401. 10.1093/jxb/erg262

Marchiori PER, Machado EC, Ribeiro RV (2014) Photosynthetic limitations imposed by self-shading in field-grown sugarcane varieties. Field Crops Research 155:30–37. 10.1016/j.fcr.2013.09.025

Marchiori PER, Ribeiro RV, da Silva L, Machado RS, Machado EC, Scarpari MS (2010) Plant growth, canopy photosynthesis and light availability in three sugarcane varieties. SugarTech 12:160–166. 10.1007/s12355-010-0031-7

Marshall B, Biscoe PV (1980) A model for C3 leaves describing the dependence of net photosynthesis on irradiance. Journal of Experimental Botany 31:29–39. 10.1093/jxb/31.1.41

Muntean L, Ona A, Berindean I, Racz I, Muntean S (2022) Maize Breeding: From Domestication to Genomic Tools. Agronomy 12:2365. 10.3390/agronomy12102365

Niinemets U, Keenan TF, Hallik L (2015) A worldwide analysis of within-canopy variations in leaf structural, chemical and physiological traits across plant functional types. New Phytologist 205:973–993. 10.1111/nph.13096

Pignon CP, Jaiswal D, McGrath JM, Long SP (2017) Loss of photosynthetic efficiency in the shade. An Achilles heel for the dense modern stands of our most productive C4 crops? J. Exp. Bot. 68:335–345. 10.1093/jxb/erw456

R Core Team (2024) R: A Language and Environment for Statistical Computing. R Foundation for Statistical Computing, Vienna, Austria.

Ray DK, Mueller ND, West PC, Foley JA (2013) Yield trends are insufficient to double global crop production by 2050. PloS one 8:e66428. 10.1371/journal.pone.0066428

Richards RA (2000) Selectable traits to increase crop photosynthesis and yield of grain crops. J. Exp. Bot. 51:447–458. 10.1093/jexbot/51.suppl_1.447

Sage RF, Pearcy RW (1987) The nitrogen use efficiency of C3 and C4 plants II. Leaf nitrogen effects on the gas exchange characteristics of Chenopodium album (L.) and Amaranthus retroflexus (L.). Plant Physiology 84:959–963. 10.1104/pp.85.2.355

Sage RF (2002) Variation in the k(cat) of Rubisco in C3 and C4 plants and some implications for photosynthetic performance at high and low temperature. Journal of Experimental Botany 53:609–620. 10.1093/jexbot/53.369.609

Sales CRG, Ribeiro RV, Hayashi AH, Marchiori PER, Silva KI, Martins MO, Silveira JAG, Silveira NM, Machado EC (2018) Flexibility of C4 decarboxylation and photosynthetic plasticity in sugarcane plants under shading. Environ. Exp. Bot. 149:34–42. 10.1016/j.envexpbot.2017.10.027

Salter WT, Merchant AM, Richards RA, Trethowan R, Buckley TN (2019) Rate of photosynthetic induction in fluctuating light varies widely among genotypes of wheat. Journal of experimental botany 70:2787–2796. 10.1093/jxb/erz100

Sarruge JR (1975) Soluções nutritivas. Summa Phytopathologica 3:231–233.

Slattery RA, Walker BJ, Weber APM, Ort DR (2018) The impacts of fluctuating light on crop performance. Plant physiology 176:990–1003. 10.1104/pp.17.01234

Thomas H, Howarth CJ (2000) Five ways to stay green. J. Exp. Bot. 51:329–337. 10.1093/jexbot/51.suppl_1.329

von Caemmerer S (2021) Updating the steady-state model of C4 photosynthesis. J. Exp. Bot. 72:6003–6017. 10.1093/jxb/erab266

von Caemmerer S, Furbank RT (2003) The C4 pathway: an effcient CO2 pump. Photosynth. Res. 77:191–207. 10.1023/A:1025830019591

Vu JCV, Allen LH Jr, Gesch RW (2006) Up-regulation of photosynthesis and sucrose metabolism enzymes in young expanding leaves of sugarcane under elevated growth CO2. Plant Sci 171:123–131. 10.1016/j.plantsci.2006.03.003

Wei S, Li X, Lu Z, Zhang H, Ye X, Zhou Y, Li J, Yan Y, Pei H, Duan F, Wang D, Chen S, Wang P, Zhang C, Shang L, Zhou Y, Yan P, Zhao M, Huang J, Bock R, Qian Q, Zhou W (2022) A transcriptional regulator that boosts grain yields and shortens the growth duration of rice. Science 377(6604), eabi8455. 10.1126/science.abi8455

Yin X, Sun Z, Struik PC, van der Putten PE, van Ieperen WIM, Harbinson J (2011) Using a biochemical C4 photosynthesis model and combined gas exchange and chlorophyll fluorescence measurements to estimate bundle-sheath conductance of maize leaves differing in age and nitrogen content. Plant, Cell and Environment 34:2183–2199. 10.1111/j.1365-3040.2011.02414.x

Yin X, van der Putten PE, Driever SM, Struik PC (2016) Temperature response of bundle-sheath conductance in maize leaves. Journal of Experimental Botany 67:2699–2714. 10.1093%2Fjxb%2Ferw104

Yoon DK, Ishiyama K, Suganami M, Tazoe Y, Watanabe M, Imaruoka S, Ogura M, Ishida H, Suzuki Y, Obara M, Mae T, Makino A (2020) Transgenic rice overproducing Rubisco exhibits increased yields with improved nitrogen-use efficiency in an experimental paddy field. Nature Food 1:134–139. 10.1038/s43016-020-0033-x

Zhu X, Long SP, Ort DR (2010) Improving photosynthetic efficiency for greater yield. Annual Review of Plant Biology 61:235–261. 10.1146/annurev-arplant-042809-112206

