## Supplementary Material for "Exploring intra-specific variation in photosynthesis of maize and sorghum"

Figure S1

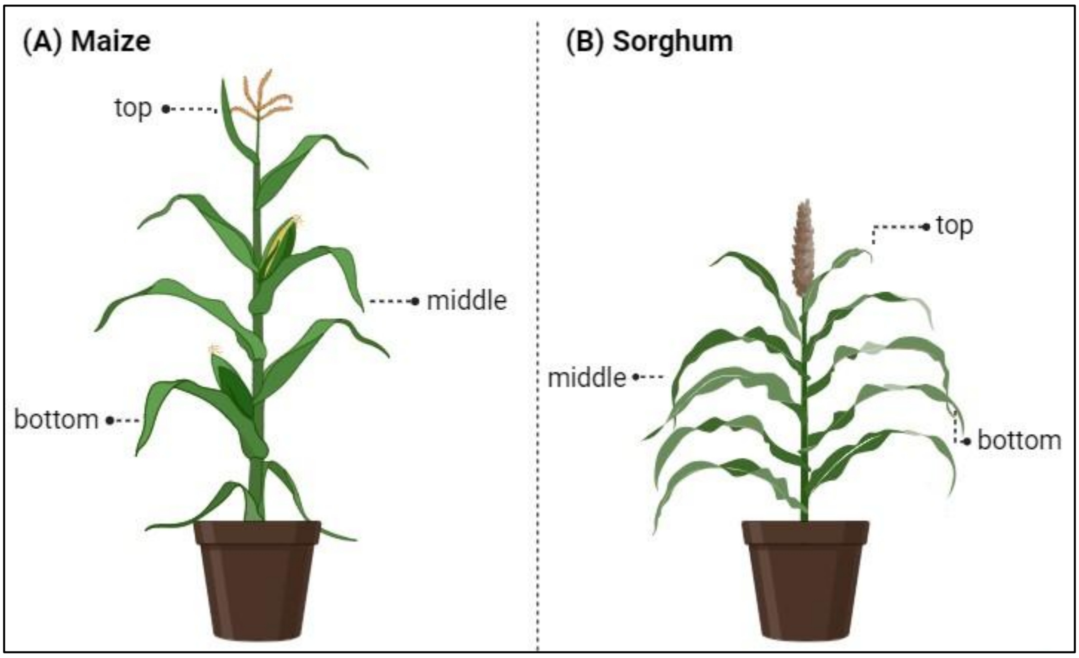


**Fig. 1.** Scheme of gas exchange evaluations in leaves on top, middle and bottom positions along the canopy of maize (in A) and sorghum (in B). Created in BioRender.com.

Figure S2

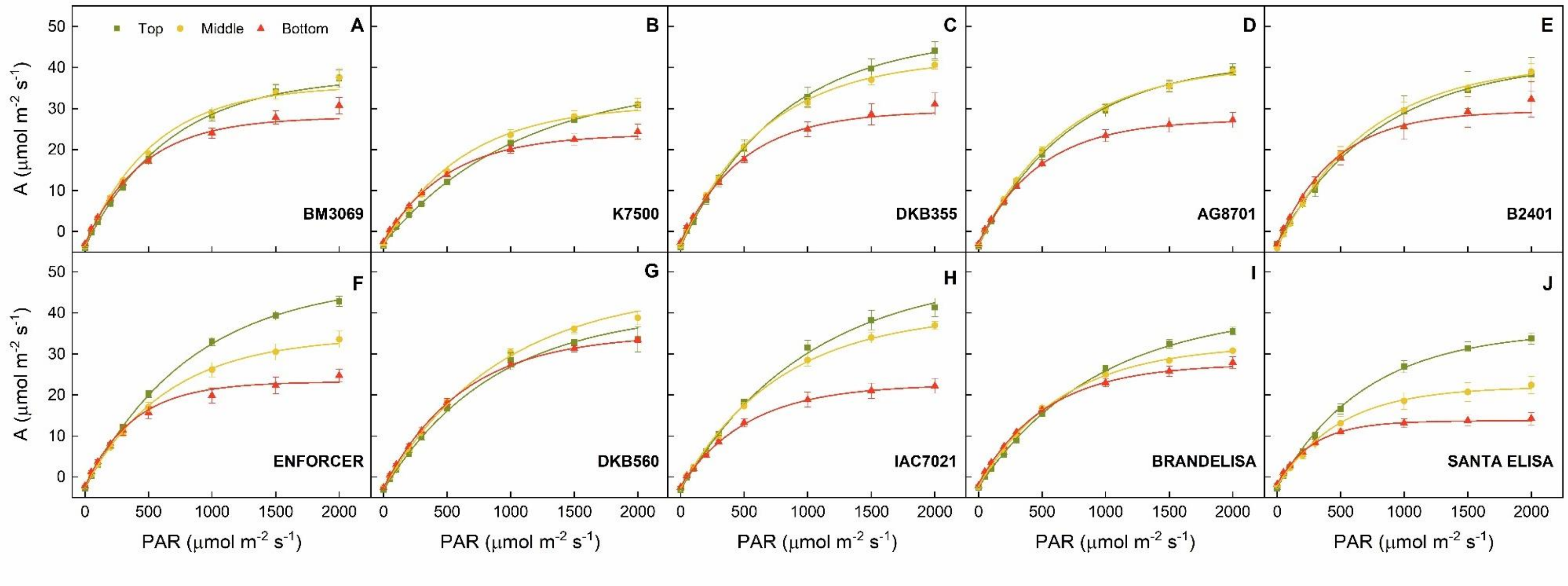


**Fig. S2**. Response curves of leaf $CO_2$ assimilation rate (*A*) to increasing light intensity (PAR) in top (green), middle (yellow) and bottom (red) leaves of maize (A-E) and sorghum (F-J) cultivars. Each symbol represents the mean±standard error (*n*=4).

Figure S3

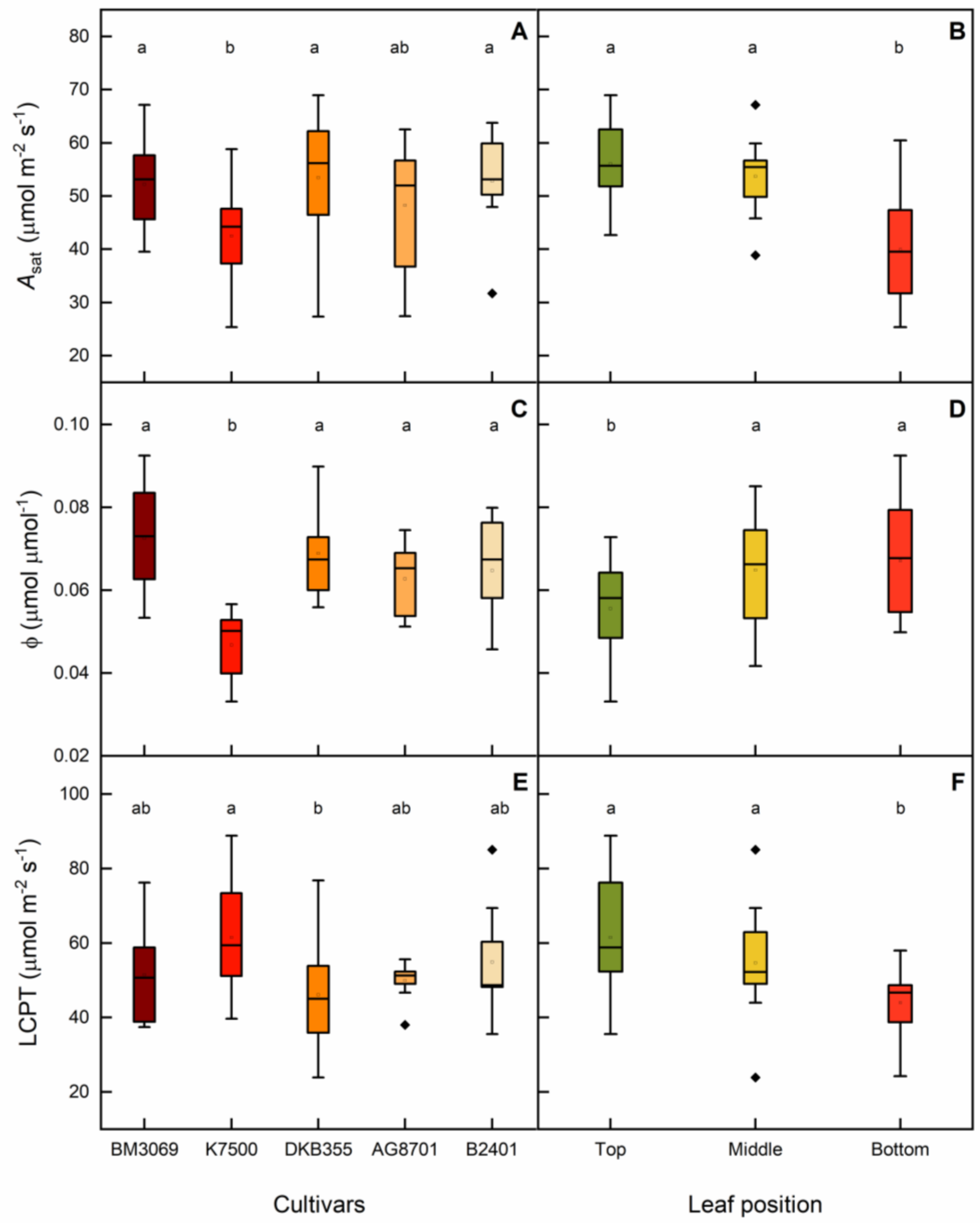


**Fig. S3.** Light saturated leaf $CO_2$ assimilation ($A_{sat}$, A and B), maximum quantum efficiency of $CO_2$ assimilation (ϕ, C and D) and light compensation point (LCPT, E and F) in five maize cultivars and three canopy layers. Different letters indicate statistical differences among cultivars (in A, C and E Tukey $p<0.05$, $n=12$), and canopy layers (in B, D and F Tukey $p<0.05$, $n=20$).

Figure S4

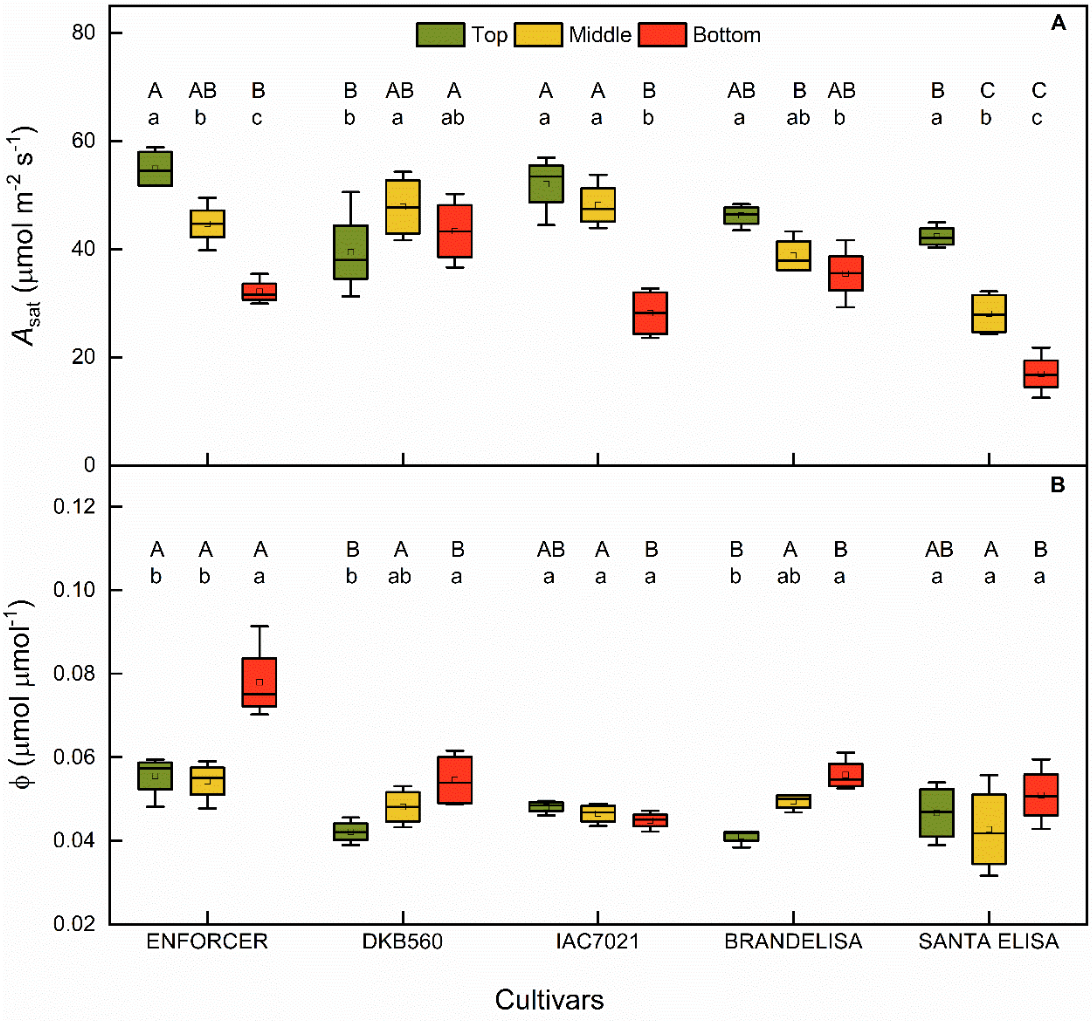


**Fig. S4.** Saturated leaf $CO_2$ assimilation ($A_{sat}$, A) and maximum quantum efficiency of $CO_2$ assimilation ($\phi$, B) in five sorghum cultivars. Different capital and lowercase letters indicate statistical differences among cultivars and canopy layers, respectively (Tukey $p$<0.05, $n$=4).

Figure S5

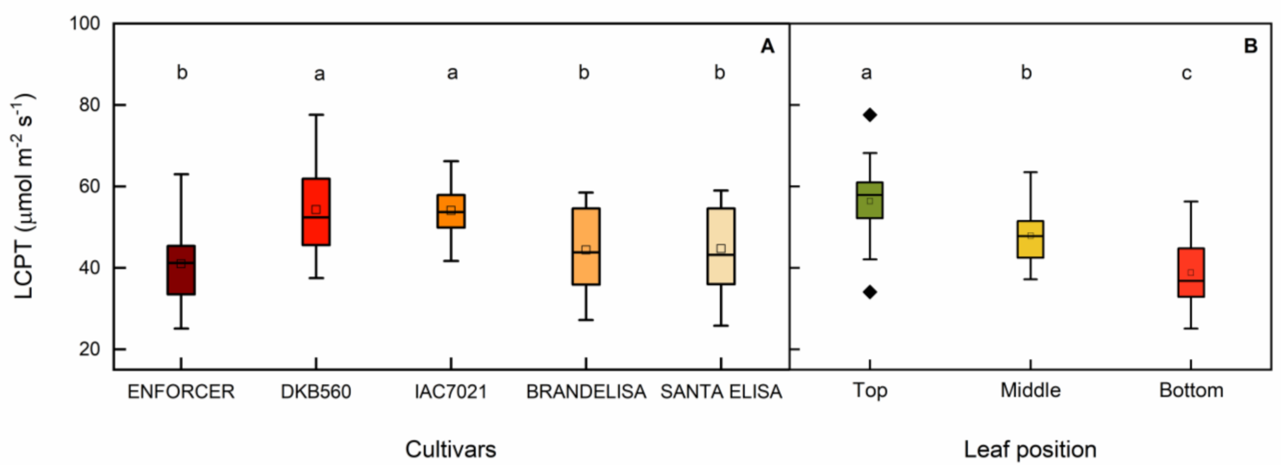


**Fig. S5.** Light compensation point (LCPT) in five sorghum cultivars and three canopy layers. Different letters indicate statistical differences among cultivars (in A, Tukey $p<0.05$, $n=12$) and canopy layers (in B, Tukey $p<0.05$, $n=20$).

Figure S6

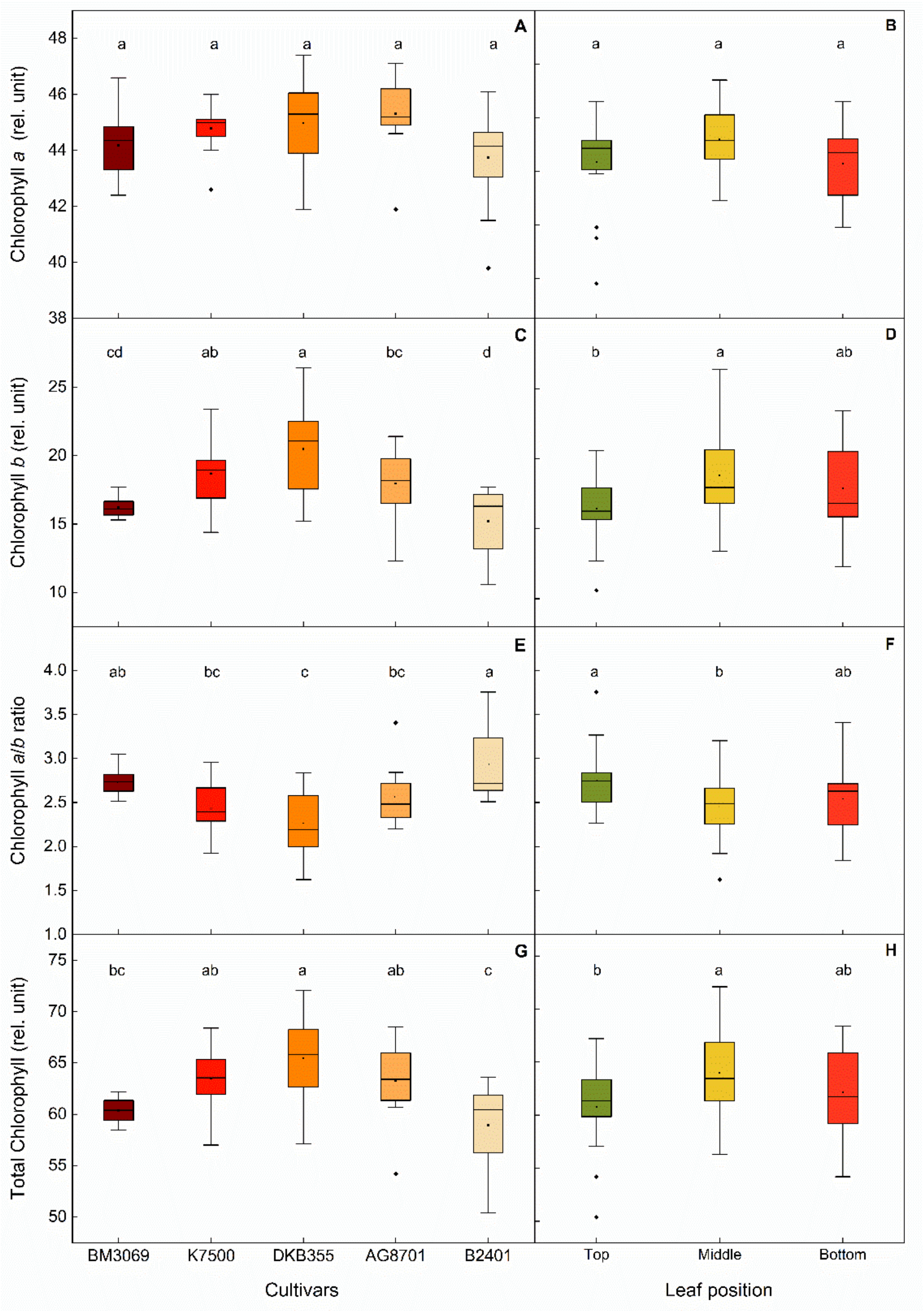


**Fig. S6.** Chlorophyll *a* (in A and B), *b* (in C and D), *a*/*b* ratio (in E and F) and total (*a*+*b*) index (in G and H) in five maize cultivars and three canopy layers. Different letters indicate statistical differences among cultivars (in A, C and E, Tukey $p<0.05$, $n=12$) and canopy layers (in B, D and F, Tukey $p<0.05$, $n=20$).

Figure S7

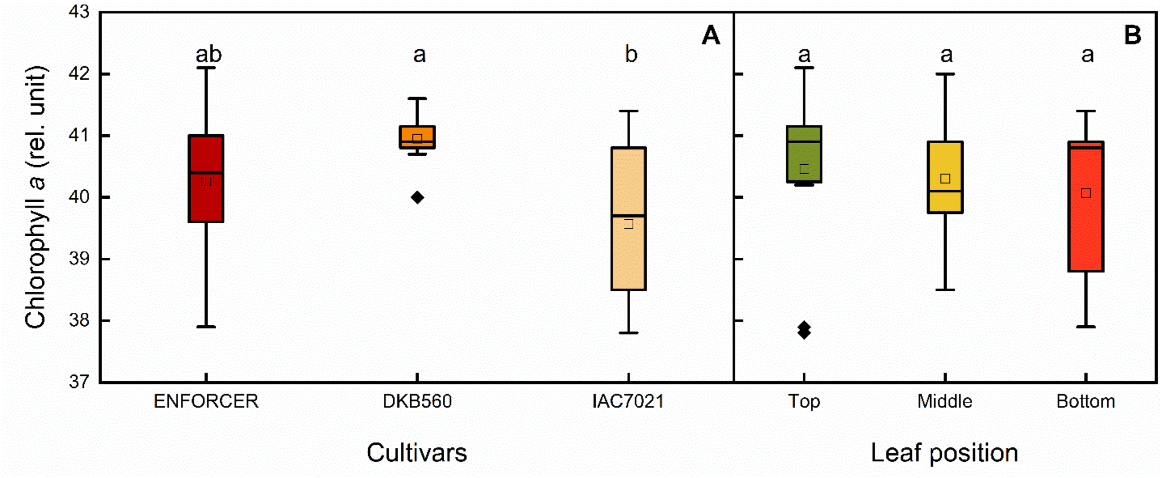


**Fig. S7.** Chlorophyll *a* index in three sorghum cultivars and three canopy layers. Different letters indicate statistical differences among cultivars (in A, Tukey $p<0.05$, $n=12$) and canopy layers (in B, Tukey $p<0.05$, $n=20$).

Figure S8

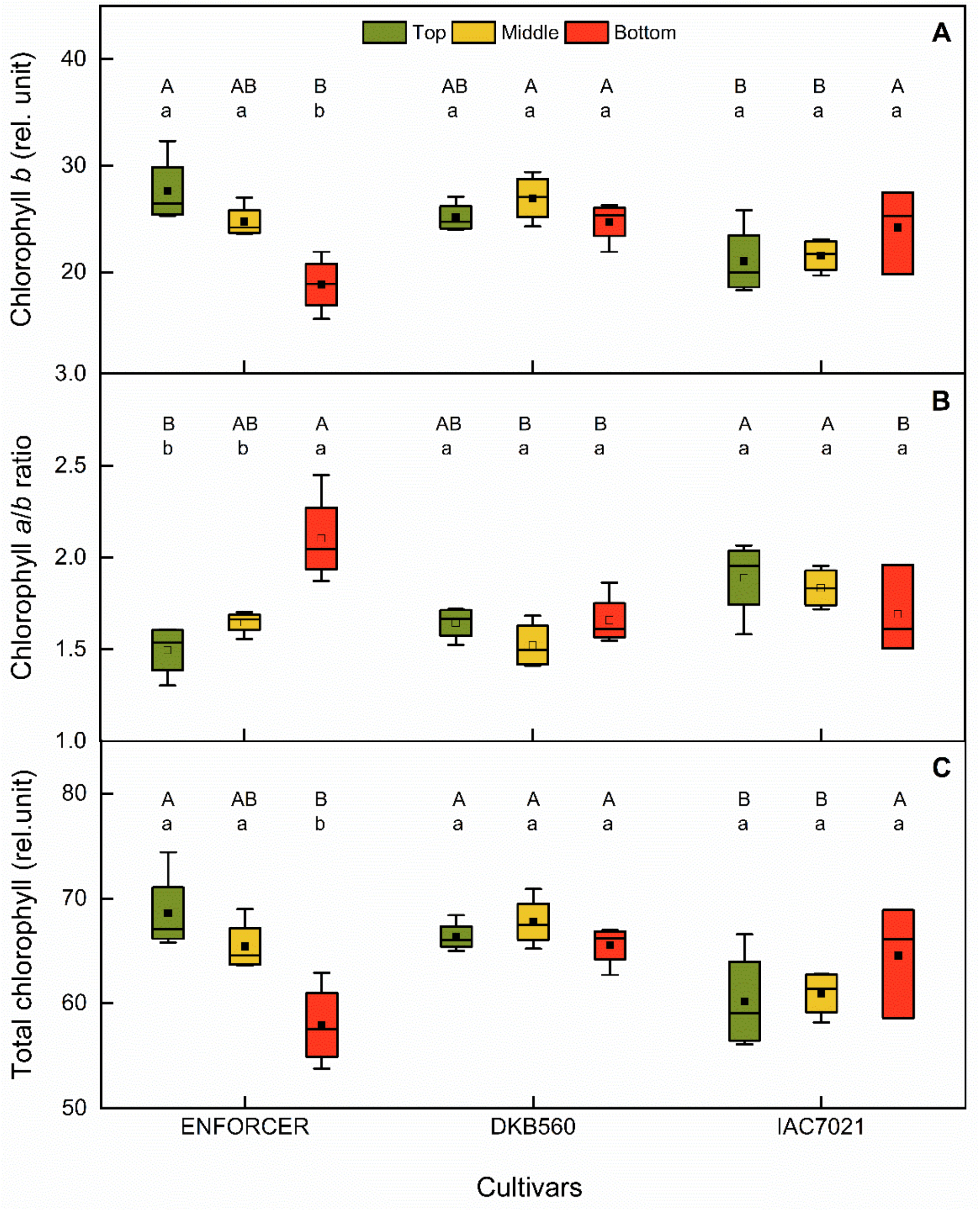


**Fig. S8.** Chlorophyll *b* (in A), *a*/*b* ratio (in B) and total (*a*+*b*) index (in C) in top (green), middle (yellow) and bottom (red) leaves of five sorghum cultivars. Different capital and lowercase letters indicate statistical differences among cultivars and canopy layers, respectively (Tukey $p<0.05$, $n=4$).

Figure S9

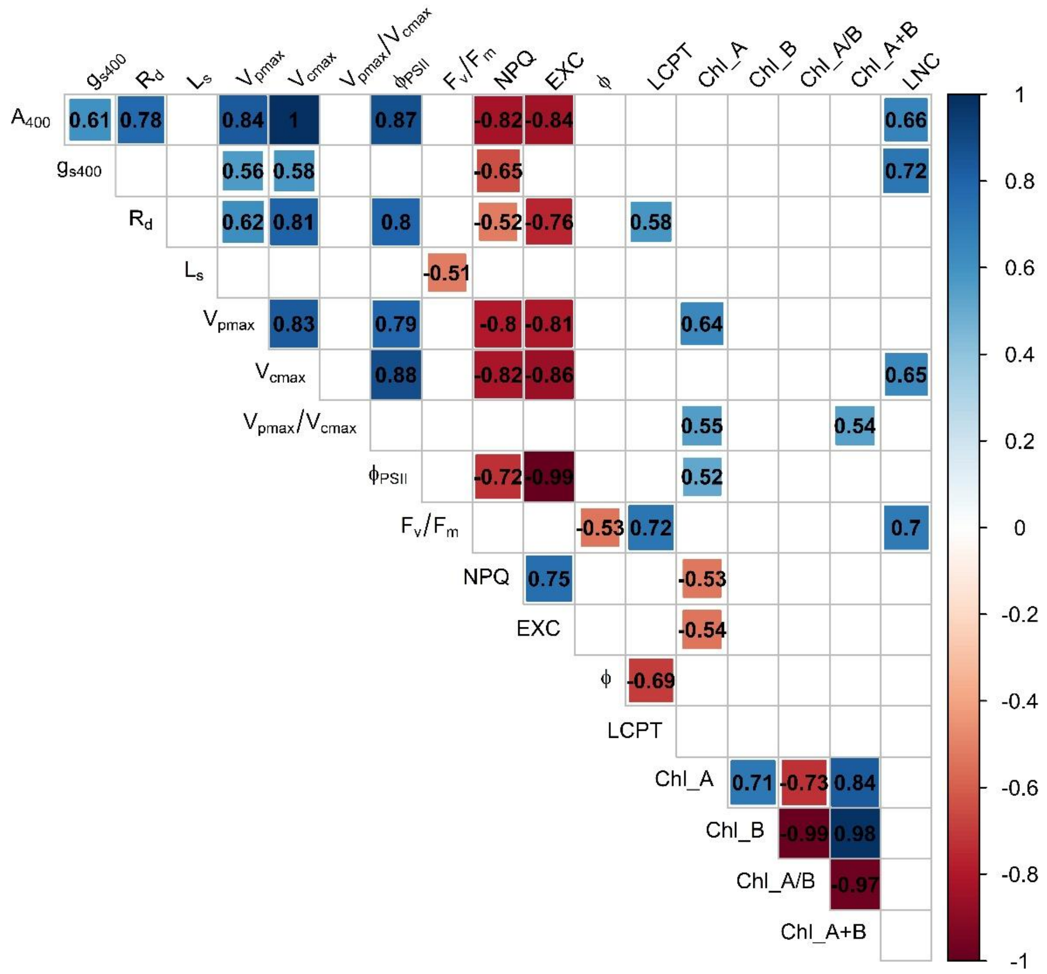


**Fig. S9.** Correlation of five maize genotypes, based on Pearson's coefficient ($p$<0.05). Leaf $CO_2$ assimilation ($A_{400}$) and stomatal conductance ($g_{s400}$) at partial air $CO_2$ pressure ($C_a$) of 400 μmol $mol^{-1}$, mitochondrial respiration in dark ($R_d$), stomatal limitation of photosynthesis ($L_S$), *in vivo* maximum carboxylation rates of PEPC ($V_{pmax}$) and Rubisco ($V_{cmax}$), $V_{pmax}$:$V_{cmax}$ ratio, effective ($\Phi_{PSII}$) and maximum ($F_v/F_m$) quantum efficiency of PSII, non-photochemical quenching (NPQ), relative excess of energy (EXC), maximum quantum efficiency of $CO_2$ assimilation (ϕ), light compensation point (LCPT), chlorophyll *a* (Chl_A), *b* (Chl_B), *a*/*b* (Chl_A/B) and *a* + *b* (Chl_A+B) indexes, and leaf nitrogen content (LNC).

Figure S10

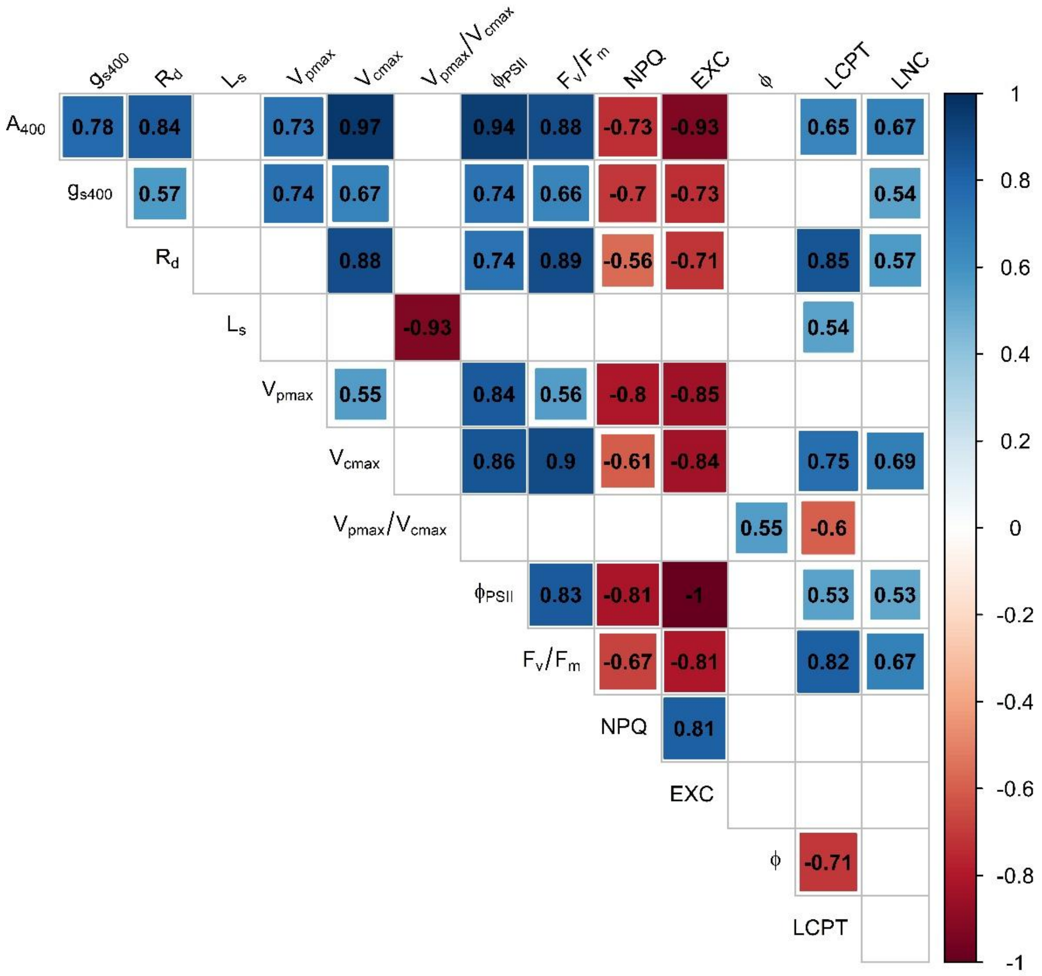


**Fig. S10.** Correlation of five sorghum genotypes, based on Pearson's coefficient ($p$<0.05). Leaf $CO_2$ assimilation ($A_{400}$) and stomatal conductance ($g_{s400}$) at partial air $CO_2$ pressure ($C_a$) of 400 μmol mol$^{-1}$, mitochondrial respiration in dark ($R_d$), stomatal limitation of photosynthesis ($L_S$), *in vivo* maximum carboxylation rates of PEPC ($V_{pmax}$) and Rubisco ($V_{cmax}$), $V_{pmax}$:$V_{cmax}$ ratio, effective ($\Phi_{PSII}$) and maximum ($F_v/F_m$) quantum efficiency of PSII, non-photochemical quenching (NPQ), relative excess of energy (EXC), maximum quantum efficiency of $CO_2$ assimilation (ϕ), light compensation point (LCPT) and leaf nitrogen content (LNC).
